# SYNTERUPTOR: mining genomic islands for non-classical specialised metabolite gene clusters

**DOI:** 10.1101/2024.01.03.573040

**Authors:** Drago Haas, Matthieu Barba, Cláudia M. Vicente, Šarká Nezbedová, Amélie Garénaux, Stéphanie Bury-Moné, Jean-Noël Lorenzi, Laurence Hôtel, Luisa Laureti, Annabelle Thibessard, Géraldine Le Goff, Jamal Ouazzani, Pierre Leblond, Bertrand Aigle, Jean-Luc Pernodet, Olivier Lespinet, Sylvie Lautru

**Author notes:** To whom correspondence should be addressed. Sylvie Lautru. Correspondence may also be addressed to Olivier Lespinet. Joint Authors (Drago Haas, Matthieu Barba, Olivier Lespinet, Sylvie Lautru). Present Address. Drago Haas, Biose Industrie, Aurillac, 15000, France. Matthieu Barba, European Bioinformatics Institute, Hinxton, CB10 1SD, UK. Cláudia M. Vincente, GenPhySE, Université de Toulouse, INRAE, ENVT, Castanet-Tolosan, France. Amélie Garénaux, Applied Medical, Rancho Santa Margarita, CA, 92688, USA. Jean-Noël Lorenzi, CNRS, Institut Jacques Monod, Paris, F-75013, France. Luisa Laureti: Team DNA Damage and Genome Instability, Cancer Research Center of Marseille (CRCM); CNRS, Aix Marseille Univ, INSERM, Institut Paoli-Calmettes, Marseille, France.

## Abstract

Microbial specialised metabolite biosynthetic gene clusters (SMBGCs) are a formidable source of natural products of pharmaceutical interest. With the multiplication of genomic data available, very efficient bioinformatic tools for automatic SMBGC detection have been developed. Nevertheless, most of these tools identify SMBGCs based on sequence similarity with enzymes typically involved in specialised metabolism and thus may miss SMBGCs coding for under characterised enzymes. Here we present SYNTERUPTOR (https://bioi2.i2bc.paris-saclay.fr/synteruptor), a program that identifies genomic islands, known to be enriched in SMBGCs, in the genomes of closely related species. With this tool, we identified a SMBGC in the genome of *Streptomyces ambofaciens* ATCC23877, undetected by earlier versions of antiSMASH, and experimentally demonstrated that it directs the biosynthesis of two metabolites, one of which was identified as sphydrofuran. SYNTERUPTOR is also a valuable resource for the delineation of individual SMBGCs within antiSMASH regions that may encompass multiple clusters, and for refining the boundaries of these SMBGCs.

## INTRODUCTION

Microbial specialised metabolites, also called natural products, are small metabolites produced by microorganisms. They exhibit a wide range of biological activities that have been extensively exploited by humans. Some of these metabolites, primarily antimicrobial and anticancer drugs, have proven essential in increasing human life expectancy (1, 2).

Others find applications in agriculture or veterinary medicine (3, 4). In natural habitats, microbial specialised metabolites play crucial roles in mediating the interactions between microorganisms and their environment. They are involved in processes such as in metal chelation, quorum sensing and mediating mutualistic, symbiotic or competitive relationships with other microbes, plants or animals (5–9). However, in many cases, the specific biological functions of these metabolites remain unknown.

Given the importance of microbial specialised metabolites, particularly as antimicrobial agents, and in the context of antibiotic resistance, a major threat to human health, there is a need to discover new metabolites and study the genes and proteins responsible for their biosynthesis. Advancements in genome sequencing have facilitated these studies by enabling the search and identification of specialised metabolite biosynthetic gene clusters (SMBGCs). The genomes of some bacterial genera, such as *Streptomyces*, *Salinispora*, and *Myxococcus*, are known to contain numerous SMBGCs, and with the increasing availability of genome sequences, bioinformatic tools are essential for efficient genome mining. Several such tools have been developed in the last 15 years and have been subject to various reviews (10–12). Most of these tools rely on classical genome mining approaches, utilizing similarity searches with known enzyme sequences typically involved in specialised metabolism, like nonribosomal peptide synthetases (NRPSs), polyketide synthases (PKSs), or terpene synthases. Prominent examples of such tools include BAGEL (13), CLUSEAN (14), NaPDoS (15), PRISM (16) and antiSMASH (17), with the latter being widely used for its ability to detect 81 different types of gene clusters. However, these sequence similarity-based tools have limitations as they can only identify gene clusters associated with already known enzyme families, leaving potential clusters involving uncharacterised genes or enzymes undetected.

To address this limitation, alternative tools have been developed using various approaches to detect SMBGCs. For example, EvoMining utilizes an evolutionary-driven genome mining strategy based on phylogeny, identifying genes in SMBGCs that may have evolved from the duplication of central metabolism genes, followed by expansion in enzyme substrate specificities (18). This approach has led to the discovery of novel biosynthetic gene clusters for arseno-organic metabolites in *Streptomyces*, which were not detected by antiSMASH. Another tool called ARTS is based on searching for paralogues of housekeeping genes. Indeed, bacteria producing antibiotics that target housekeeping proteins may produce a resistant version of the target protein, which can often be encoded in SMBGCs (19). Here we propose another type of approach, based on the detection of genomic islands in the genomes of closely related species.

Genomic islands are regions present in the genomes of some closely related species but absent in others and are typically acquired through horizontal gene transfer. These islands are surrounded by syntenic genomic regions, enabling their detection. The first identified genomic islands were pathogenic islands, but since then, various functional types have been discovered, including symbiosis islands, secretion islands, antibiotic resistance islands or metabolic islands (20, 21). Given that SMBGCs are known to be exchanged through horizontal gene transfer (22–24), they are likely to be found within genomic islands. Indeed, several studies over the past decade have shown that genomic islands are frequently enriched in SMBGCs, particularly in prolific producers of specialised metabolites like Actinomycetota. For instance, in studies focusing on the marine Actinomycetota *Salinispora*, it has been shown that SMBGCs are frequently found within genomic islands that are well-conserved among *Salinispora* strains (22, 25, 26).

Based on the enrichment of SMBGCs in genomic islands, we hypothesized that the identification of genomic islands in the genome of a bacterial species could serve as a promising starting point for isolating SMBGCS, including those not detectable by classical tools such as antiSMASH. To address this hypothesis, we developed a bioinformatic tool named SYNTERUPTOR, which identifies genomic islands in a given genome by comparing its genomic sequence with those of closely related species. While bioinformatics tools for genomic island identification have been developed (27–29), SYNTERUPTOR was designed and is focused on identifying SMBGC-containing genomic islands. Indeed, the content of the identified genomic islands is analysed using in particular SMBGC related or specific criteria. Thus, the tool offers the user assistance in selecting specific genomic islands susceptible to contain SMBGCs. SYNTERUPTOR can be used alongside existing tools like antiSMASH or ARTS to enhance the mining of bacterial genomes and the discovery of atypical SMBGCs. Using this tool, we identified a SMBGC in the genome of *Streptomyces ambofaciens* ATCC23877 and experimentally demonstrated that it directs the biosynthesis of two metabolites, one of which was identified as sphydrofuran. This demonstration underscores the utility of SYNTERUPTOR in genome mining and the discovery of unique SMBGCs.

### MATERIAL AND METHODS

### The SYNTERUPTOR pipeline

As an input, the SYNTERUPTOR pipeline requires a dataset consisting of genome files selected by the user from species that are related enough to possess synteny blocks. It proceeds by performing pairwise comparisons between all Coding DNA Sequences (CDSs) amino acid sequences to identify orthologs. Subsequently, it constructs synteny blocks and detects any instances of synteny breaks.

### Synteny block detection

First, the CDS amino acid sequences in all genomes are compared with each other using Blastp (30, 31). The orthologs between each pair of genomes are then computed using BRH (Best Reciprocal Hits), which is well adapted for closely related genomes (32). When several best hits are detected, synteny is used to assign orthology. Two sequences are considered orthologs if they are both preceded or followed by another identified ortholog pair in both genomes. If there are multiple best hits remaining for a CDS, no ortholog pair is retained.

To build synteny blocks, we used the method developed for Syntebase (33): the data are introduced in a database and the synteny blocks are computed using SQL queries. Once the database is constructed with the genes coordinates and the list of orthologs, all consecutive orthologs in all pairs of genomes are grouped in pairs, allowing for overlaps between them. In order to tolerate potential annotation and orthology detection errors, and because we do not expect to find very small breaks informative, we allowed small gaps in synteny blocks.

Specifically, orthologs within a pair can be separated by up to two non-orthologous CDSs. Subsequently, all pairs are then expanded and merged to form blocks by aggregating any overlapping pairs of orthologs. Importantly, all orthologs in the final blocks are in the same order in both genomes. Finally, the blocks are numbered based on their order along each respective genome.

### Synteny break detection

Synteny breaks are defined as genomic regions that occur between two consecutive synteny blocks in two genomes, with the two blocks on the same strand. Synteny breaks are therefore detected when two consecutive blocks in one genome are also consecutive in the other compared genome.

In order to tolerate some orthology detection errors, we also allowed the presence of a small number of blocks within the breaks (as long as they are not consecutive in both genomes).

### Break analysis

We then analyse all the breaks to determine their gene content, based on the genome annotation and comparing it with a list of keywords of interest related to mobility (such as: insertion, mobile element, integrase, transposase) or functions of interest (such as resistance, specialised metabolism biosynthetic enzymes). Additionally, we take into account other relevant information for each break, such as the presence of a tRNA within the break in both genomes, which is a hint for potential hotspots of horizontal gene transfer.

Furthermore, we retain the paralogs obtained from running a blastp search of all the proteins encoded in a genome against themselves. This information provides additional insights into the genomic organization and potential gene duplications within each genome.

### Break viewer

In order to easily explore the data within a SYNTERUPTOR database generated from a given set of genomes, we developed a user-friendly web interface that can be used in any modern web-browser. The viewer has three primary components: the dotplot, the ranking page and the break viewer.

*Dotplot.* The dotplot provides a graphical representation of the synteny relationships between the genomes, allowing users to visually identify and analyse the patterns of synteny blocks and breaks. Between any two genomes in a given database, the dotplot depicts the orthologs as grey circles, synteny blocks as black lines, and breaks as red rectangles (exact coordinates) and circles (highlighting the breaks position at any zoom level). This representation allows users to observe and analyse the relationships between genomes by highlighting the presence of orthologs, the boundaries of synteny blocks, and the locations of breaks. In addition, the GOC (Gene Order Conservation) profile, which corresponds to the number of adjacent orthologs divided by the number of CDS in a sliding window measuring 3% of the genome size, is shown along each genome (34).

The dotplot graph is interactive and enhances user exploration by allowing zooming and filtering to customize the view based on the size of breaks. This flexibility enables users to explore a broad range of synteny disruptions. Each break displayed in the dotplot includes links to the corresponding break viewer. This feature allows users to conveniently access detailed information and further analyse the specific characteristics, gene content, and other relevant data associated with each break of interest.

*Ranking page.* The ranking page presents a comprehensive list of all breaks between two genomes within the SYNTERUPTOR database. Each break is accompanied by a range of properties that can be utilised for ranking purposes. These properties include the size of the breaks, the presence or absence of tRNAs, and any other relevant characteristics. To facilitate a customized and focused analysis, the ranking page offers filter options. One such filter allows users to specify a minimum size requirement (number of CDSs) for breaks in each genome. By setting this filter, users can narrow down the list to display only breaks that meet their desired size criteria. Another option allows the user to assign different weights to parameters (Table S1) for ranking identified genomic islands.

*Break viewer.* The break viewer provides a detailed visualization of all the genes within a given break between two genomes. It aligns both genomes based on their flanking synteny blocks, allowing for a clear comparison and examination of the gene content within the synteny break i.e. within the genomic island.

### Software availability

The SYNTERUPTOR Pipeline was implemented as a collection of Perl and Python scripts utilising the BioPerl and BioPython packages, along with additional bash scripts. To handle the database management, Sqlite3 was employed due to its ability to store data in a single, easily shareable file.

The SYNTERUPTOR Viewer was developed as a collection of PHP web pages that utilize html5 and JavaScript. The viewer leverages libraries such as jQuery and d3.js for handling graphical data and enhancing interactivity. To run the SYNTERUPTOR Viewer, a LAMP (Linux, Apache, MySQL, PHP) environment is required.

The website, https://bioi2.i2bc.paris-saclay.fr/synteruptor, provides access to the viewer, allowing users to explore the database used in this paper. Additionally, users have the option to create their own SYNTERUPTOR Database through the provided interface. The maximum number of genome sequences that can be uploaded through the webpage is 20.

The SYNTERUPTOR source code is available in Zenodo [10.5281/zenodo.10424133], at [DOI 10.5281/zenodo.10424123] at https://github.com/i2bc/synteruptor and https://github.com/i2bc/synteruptor_web.

### Strains, plasmids and culture conditions

The strains, plasmids and cosmids used in this study are summarized in Supplementary Table S2. The construction of the mutant strains OSC4 and OSC416 is described in the supplementary data. *Escherichia coli* strains were grown in LB medium at 37°C with appropriate antibiotics when necessary. *Streptomyces* strains were grown at 30°C on solid SFM medium (Soya Flour Mannitol) for genetic manipulation and spore stock preparations (35). For the production of sphydrofuran, we used a modified R2YE medium (R2YEm) with a reduced saccharose concentration (75 mM, ¼ of the standard concentration (35)), and strains were grown in liquid culture for five days at 30°C under agitation. For the OSMAC (One Strain Many Compounds) approach, the *Streptomyces* mutant strains OCS4 and OSC416 were grown under agitation in different culture media, including R2, R2YE, R2YEm, HT and MP5 (35, 36) during 5 days at 30°C before submitting the supernatant to HPLC analysis.

### Preparation and DNA manipulation

*E. coli* transformations and *E. coli* / *Streptomyces* conjugations were performed under standard conditions (35, 37). Taq polymerase (Qiagen) was used for DNA amplifications. DNA fragments and PCR products were purified using the NucleoSpin Extract II (Macherey-Nagel). All oligonucleotides used in this work are listed in Supplementary Table S3.

### Heterologous expression of the genomic island 14594 (0762fd) in *Streptomyces coelicolor* M1154

A cosmid library of *S. ambofaciens* genomic DNA was constructed using the pWED4 cosmid. Total DNA was partially digested with Sau3AI and fragments of 35 to 45 kb were ligated with BamHI digested pWED4. The construction of the library was achieved as previously described (38). An internal fragment of SAM23877_3931 was amplified using the oligonucleotides mmyD-F and mmyD-R to generate a radiolabeled probe. This probe was used to screen the cosmid library as previously described (38). Twelve clones were isolated and their cosmid DNA (pSLM001 to pSLM012) was extracted. The presence of the SAM23877_3931 gene was verified by PCR with the oligonucleotides mmyD-F and mmyD-R. To determine which cosmids contained the complete genomic island, the presence of SAM23877_3916 (primers SL26 and SL27) and of SAM23877_3940 (primers SL24 and SL25), located just upstream and downstream of GI #14594, was searched by PCR in the twelve cosmids. Among the four cosmids that were found to contain the complete genomic island, the extremities of the inserts of two of them, pSLM003 and pSLM010 were determined by sequencing. The pSLM003 was introduced by intergeneric conjugation from *E. coli* ET12567/pUZ8002 into *S. coelicolor* M1154. Exconjugants were selected on apramycine (50 µg/mL). The resulting strains were named SPFSH001.

### Chemical purification of M1 (sphydrofuran)

One litre of culture supernatant of SPFSH001 grown in R2YEm was evaporated to dryness under reduced pressure and the residue obtained was extracted with methanol. The methanol extract was subjected to flash chromatography on a Combiflash Companion using a Redisep 40 g silica column, using a mixture of CH_2_Cl_2_/MeOH as eluent. The fractions containing M1 were identified by TLC analysis and pooled. M1 purification was achieved by prep RP-HPLC using a Waters-Alliance 2695 HPLC instrument with PDA and ELS detection equipped with a prep RP-HPLC (Sunfire Prep C18 5μm, 10 x 250 mm) and a linear H_2_O/CH_3_CN gradient supplemented with 0.1% of formic acid (100-0% to 0-100%). After concentrating *in vacuo*, M1 (19 mg) was obtained as a yellowish oil.

### HPLC analyses

After five days of culture at 30°C in R2YEm, culture supernatants were filtered through Mini-UniPrep filters (GE Healthcare Life Science) and analysed on an Atlantis dC18 column (250 mm x 4.6 mm, 5 µm, column temperature 30°C) using an Agilent 1200 HPLC instrument equipped with a quaternary pump. Samples were eluted with 0.1% HCOOH in H_2_O (solvent A) / 0.1% HCOOH in CH_3_CN (solvent B) (95:5) at 1 ml / min for 5 min followed by a gradient to 71:29 A/B over 15 min.

### HPLC-MS analyses

LC-MS experiments were performed using a Waters-Micromass ZQ2000 simple-stage quadrupole mass spectrometer equipped with an ESI (electrospray ionization) interface coupled to an Alliance Waters 2695 HPLC instrument with PDA and ELS detection.

### HR-ESI-MS analysis

HR-ESI-MS analysis was conducted using a Waters-Micromass mass spectrometer equipped with an ESI-TOF (electrospray-time-of-flight).

### NMR

NMR experiments were performed using a Bruker Advance 500 MHz spectrometer. The spectra were acquired in MeOD (δ_H_ 3.31 ppm; δ_C_ 49.9 ppm) at room temperature.

## RESULTS AND DISCUSSION

### Design of SYNTERUPTOR

We designed the SYNTERUPTOR program to compare the chromosomal sequences of closely related bacterial species (i.e. chromosomes exhibiting a high degree of synteny) and to identify and analyse the genomic islands present in each respective chromosome. The specifications of the programme were as follows.

(i) Comparative Analysis and Visualization: SYNTERUPTOR should detect the genomic islands present in the genomes of two closely related bacterial species. Furthermore, the software should support the comparison of multiple genomic sequences simultaneously and provide a comprehensive comparative analysis and interactive graphical visualization of the genomic islands across different genomes.
(ii) Analysis of the identified genomic islands: the program should perform an analysis of the genomic island content to guide the search for those with a potential SMBGC. For this purpose, we identified five criteria: i) the number of CDSs they contain, ii) the number of CDSs without ortholog in the second genome, iii) the number of CDSs with paralogs in the considered genome, iv) the presence of tRNA genes within or in the immediate vicinity of the genomic island, and (v) the difference of GC ratio between the genomic island and the complete genomic sequence. Indeed, genes located in a genomic island with orthologs somewhere in the second genome might not be part of a SMBGC. Genes that have paralogs within the considered genome can provide a potential hint towards the presence of a SMBGC, especially when these paralogs are related to enzymes involved in primary metabolism or when they represent duplicated housekeeping genes. For example, the gene cluster directing the biosynthesis of Calcium-Dependent Antibiotics (CDA) in *S. coelicolor* contains paralogs of four genes involved in the biosynthesis of tryptophan, one of the precursors of CDA (39). More importantly, the work of Cruz-Morales *et al.* has shown that enzymes from primary metabolism can be repurposed for specialised metabolism and that the identification of these enzymes can lead to the discovery of new metabolites (16). Finally, one of the resistance mechanisms to antibiotics targeting essential cellular processes is the presence, often within the SMBGC, of a gene coding for a resistant version of the target protein. This is for example the case of the antibiotics novobiocin, targeting the gyrase GyrB, and salinosporamide A, targeting the proteasome, whose SMBGCs encode a novobiocin-resistant GyrB and a salinosporamide A-resistant proteasome β-subunit, respectively (40, 41). The last two criteria, the presence of tRNA genes within or in the immediate vicinity of the genomic island, and the difference of GC ratio between the genomic island and the complete genomic sequence, were chosen as they are relevant of potential horizontal gene transfer events. Indeed, several studies have highlighted the role of these events in the acquisition of SMBGCs by microorganisms (20, 21, 42). A difference of GC ratio is one of the ways to identify a region acquired by horizontal gene transfer and tRNA genes are frequent sites for the integration of mobile elements that largely drive horizontal gene transfer in some bacteria genera such as *Streptomyces* (43, 44).
(iii) Graphical overview and tabular view: SYNTERUPTOR should provide a graphical overview of the genomic regions encompassing the identified genomic islands. Additionally, it should offer a detailed tabular view of the content within each genomic island. This tabular view should include comprehensive information about the genes, their functional annotations, and any other relevant data associated with the genomic islands.
(iv) To facilitate a customized and focused analysis, it should be possible to rank the obtained genomic islands by assigning a user-selected coefficient to the parameters defined in Table S1, such as the presence of tRNA gene adjacent to the genomic island or the number of CDSs in the genomic islands of one or the other genome.

### Pipeline for the detection of genomic islands in the chromosome of closely related species

An outline of the pipeline used to determine and analyse the genomic islands is presented in Figure 1. The first step consists of pairwise comparison of the sequences of all the predicted proteins encoded in the studied chromosomes, and in the determination of the ortholog pairs using the Best Reciprocal Hits (BRH) method (39). Pairs of sequences are retained if their E-value is smaller than 10E-10, their sequence identity is over 40% and if the sequences align to at least 40% of the length of the smaller of the two CDS. While other methods such as Syntebase (29) or RGP finder (30, 31) use 80% of the length, we observed that using 40% yielded slightly more results, possibly because blast is a local alignment program that does not extend alignments over conserved regions with low conservation. Synteny blocks are then constructed by clustering all consecutive orthologs as described in the Material and Methods section. In a given genome, synteny breaks are defined as the genomic regions located between two consecutive synteny blocks in the two compared genomes (genomic islands). In addition, for each genome, a GOC (Gene Order Conservation) profile calculated along the length of the genome is displayed, highlighting regions of conservation between the two genomes or, conversely, regions of synteny break (see Material and methods for more details). The GOC profile complements the dotplot and enables the genome-wide visualization of synteny regions.

**Figure 1.**
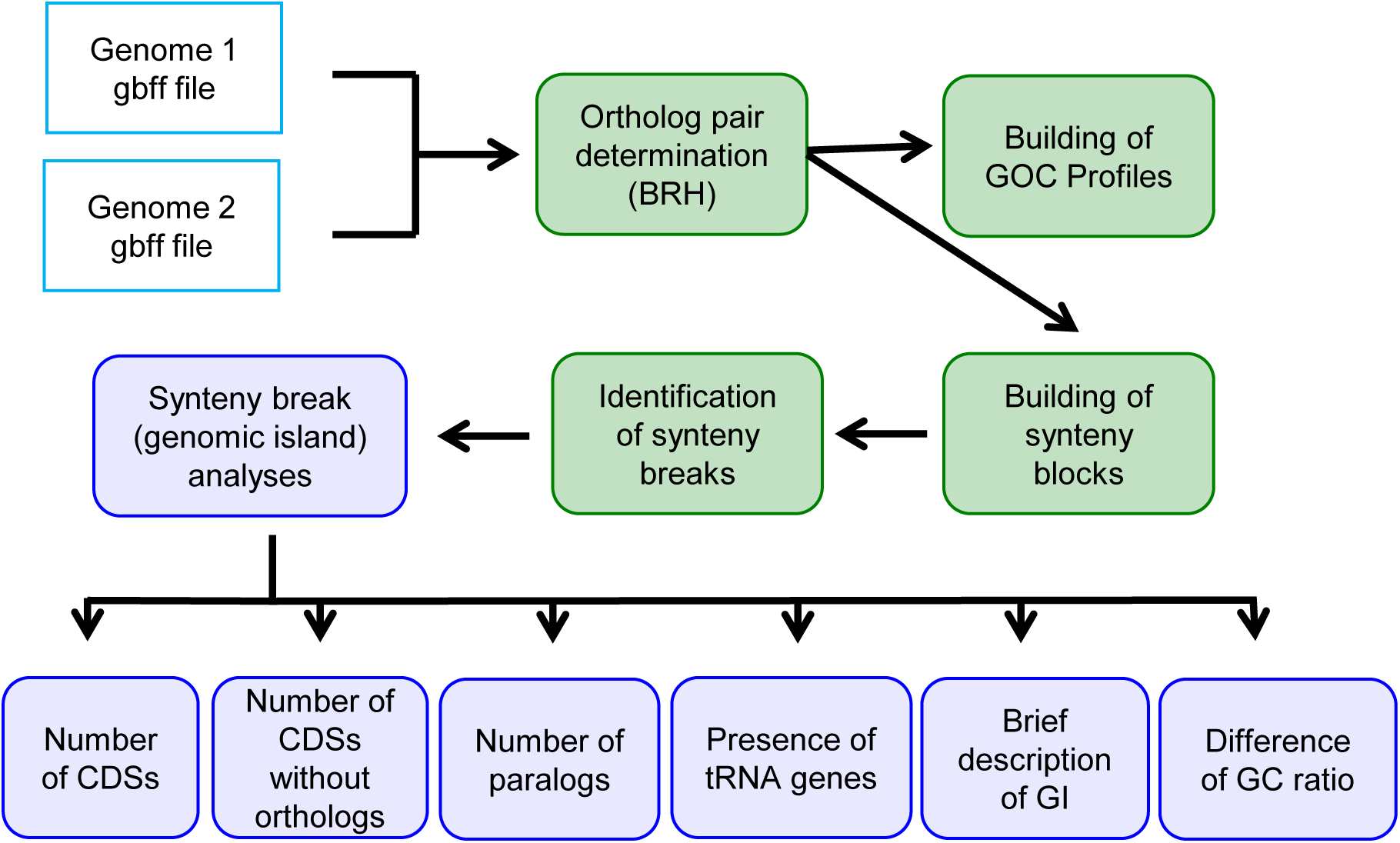
Outline of the pipeline used for the identification of the genomic islands. After gene prediction, ortholog pairs are determined. Synteny blocks are constructed by clustering all consecutive orthologs and synteny breaks (genomic islands) between two consecutive synteny blocks are identified. The genomic islands are then analysed to determine a number of parameters such as number of CDSs, number of CDSs without ortholog. BRH is for Best reciprocal BLAST hits.

The genomic islands identified are next analysed to determine (i) the number of CDSs they contain, (ii) the number of CDSs without ortholog in the second genome, (iii) the number of CDSs with paralog(s) in the considered genome, (iv) the presence of tRNA genes within or in the immediate vicinity of the genomic island, and (v) the difference in GC ratio between the genomic island and the complete genomic sequence. The results of the identification of the genomic islands, each possessing a unique identifier, and of their analyses are visualized using dotplot graphs or tables, as illustrated with the example described below.

### Detection of the genomic islands present in the genomes of *Streptomyces ambofaciens* ATCC23877 and *Streptomyces coelicolor* A(3)2

To demonstrate the functionality of SYNTERUPTOR, we compared the genomes of *Streptomyces ambofaciens* ATCC23877 and *Streptomyces coelicolor* A(3)2. We selected these species due to their relatively close phylogenetic relationship, with an average nucleotide identity using Blast (ANIb) of 86.8% on 57.3% of the sequence, as calculated using the JSpeciesWS web server (40). Moreover, the specialised metabolism of both strains has been extensively studied and is well characterised (41, 42).

The genomes of *S. ambofaciens* ATCC23877 and *S. coelicolor* A(3)2 contain 7759 and 7909 CDSs, respectively. A total number of 5366 ortholog pairs were identified. The results of the construction of the synteny blocks and of the determination of the genomic islands in both genomes are depicted in Figure 2.

**Figure 2.**
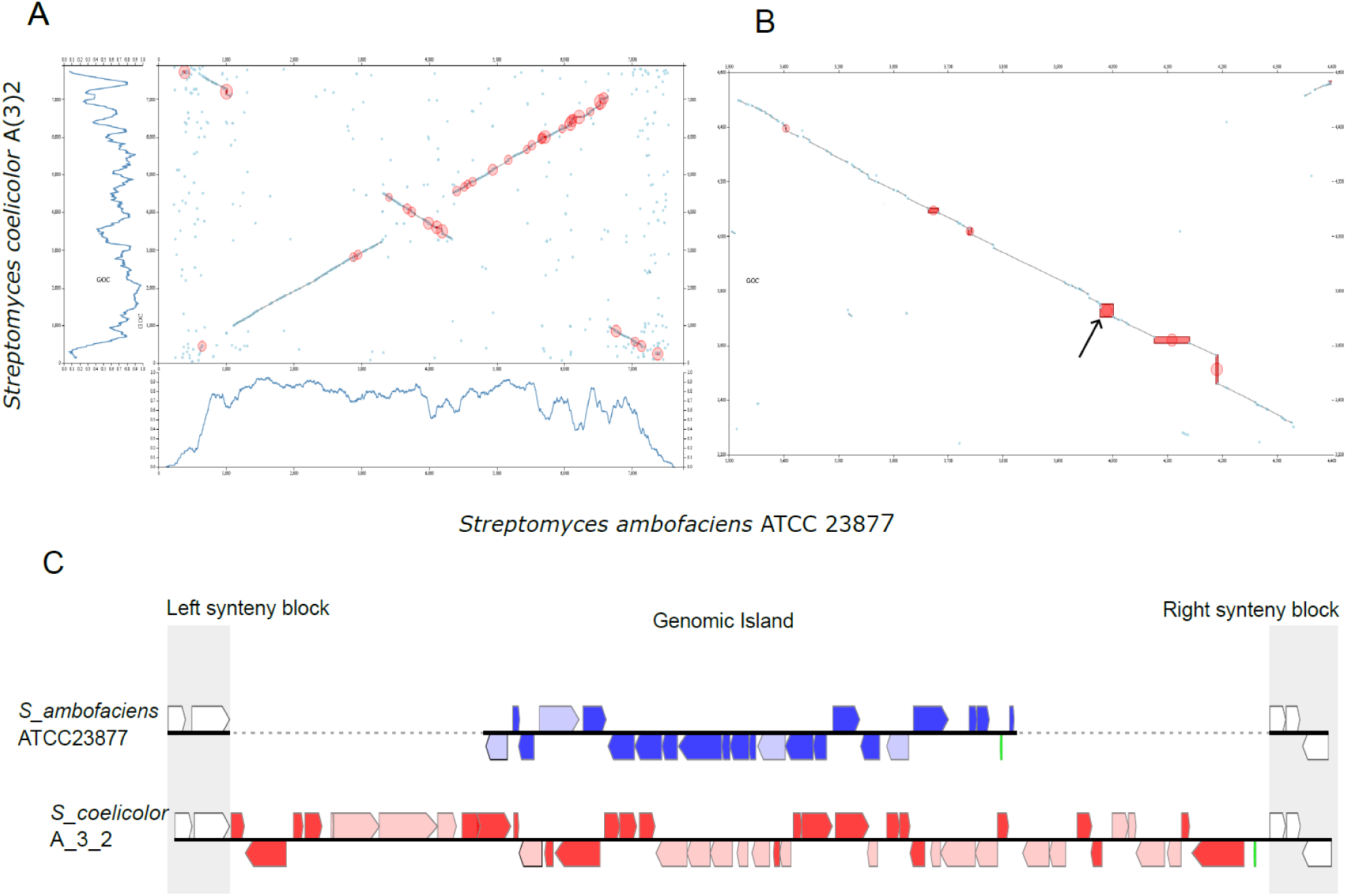
Graphical representation of the genomic islands identified by comparing the genomes of *S. ambofaciens* ATCC 23877 and *S. coelicolor* A(3)2. Only genomic islands of at least 15 CDSs found in at least one genome are shown. (A) Dotplot showing ortholog pairs and synteny blocks, with the genomic islands circled in red. GOC profiles are displayed below or on the left of the dotplot. The scale on the x-axis indicates the gene position in gene number, not in bp. (B) Close-up of the central genomic regions. The red boxes represent the GIs observed either in the genome of *S. coelicolor* (vertical bars) or in the genome of *S. ambofaciens* (horizontal bars) The arrow indicates the genomic island visualized in C. (C) Visualization of the gene content and organization of the genomic island 0762fd (#819). Genes in dark colours do not possess orthologs in the other genome, genes in light colour do. Green bars represent tRNA genes. Left and right blocks represent synteny regions. This figure represents snapshots of the web interface.

In Figure 2A, a dotplot displays the synteny blocks, with genomic islands highlighted in red circles. GOC profiles are shown on the left and bottom sides of the dotplot, allowing the visualization of highly syntenic regions. The program allows users to zoom into specific regions (Figure 2B). The content of each genomic island can be visualised as presented in Figure 2C. It is followed by a concise table summarizing the characteristics of the genomic islands. These include the number of CDSs with or without orthologs in the other genome, the number of CDSs with paralogs in the considered genome, the difference of the GC ratio between the island and the complete genome sequence, and a brief description of the functions encoded by the genes in the genomic islands such as regulation, resistance or transport. Additionally, six tables provide a brief description of each gene within the left and right synteny blocks (e.g., see Figure 3 for a left synteny block) and within the genomic islands of both species. For each gene, the sequence of the protein product can be blasted against the sequences of the group of the locally compared chromosomes or against the NCBI databases. The tables also indicate the gene ID and whether there exists an ortholog in the second genome or a paralog within the considered genome.

**Figure 3.**
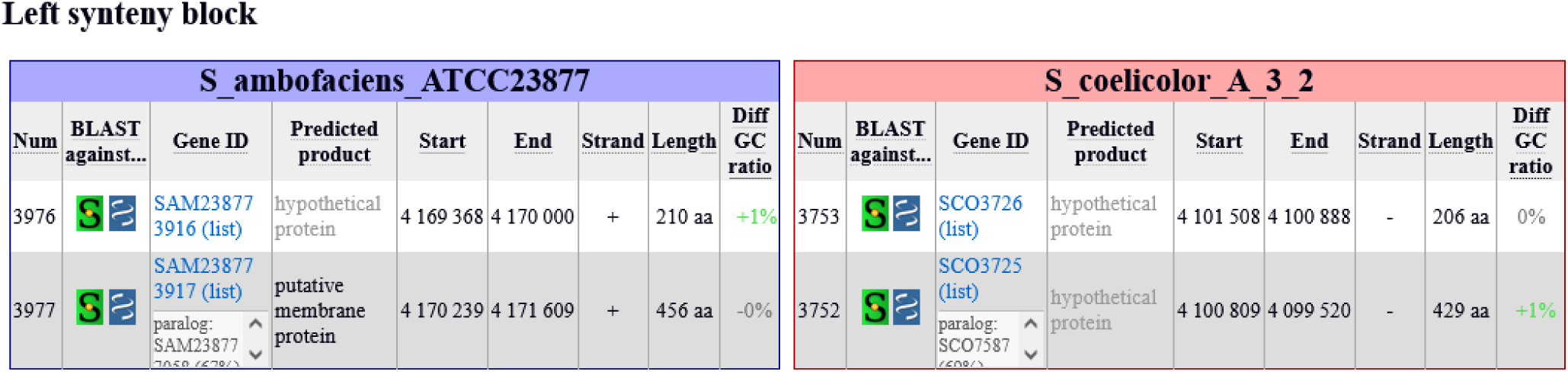
Tables describing the genes located in the left synteny blocks of the 0762fd genomic island. The gene ID, the position in the chromosome, the difference of the GC ratio between the gene and the complete genome sequence, as well as the predicted function and length of the gene product are presented. In this example, two genes, one in each genome (SAM23877_3917 and SCO3725), have a paralog in their respective chromosome. Num: CDS number. These tables represent a snapshot of the web interface.

### Setting the minimal size of the genomic islands detected

The SYNTERUPTOR program has a minimum detection size for a synteny break of three genes (CDSs) in at least one genome. Thus, there may not be any synteny break in the second genome at this position and if any, the break may contain only one or two CDSs. In our specific example, SYNTERUPTOR identified a total of 83 genomic islands in the *S. ambofaciens* ATCC23877 genome, ranging in size from 3 to 111 genes, and 124 genomic islands in the *S. coelicolor* A(3)2 genome, containing 3 to 151 genes. To allow users to focus on large genomic islands, the program offers the option to set a minimum size (in CDS number, larger than three) for the displayed genomic islands. Figure 4 illustrates the relationship between the number of genomic islands and the minimum number of genes/CDSs in those genomic islands (*S. ambofaciens* ATCC 23877 genome). The number of genomic islands decreases exponentially as the minimum size requirement for genomic islands is increased. It should be noted that in this figure, genomic islands containing zero, one or two CDSs can be visualised. They correspond to regions of the analysed genome for which a genomic island (of three or more CDSs) is present in the compared genome.

**Figure 4.**
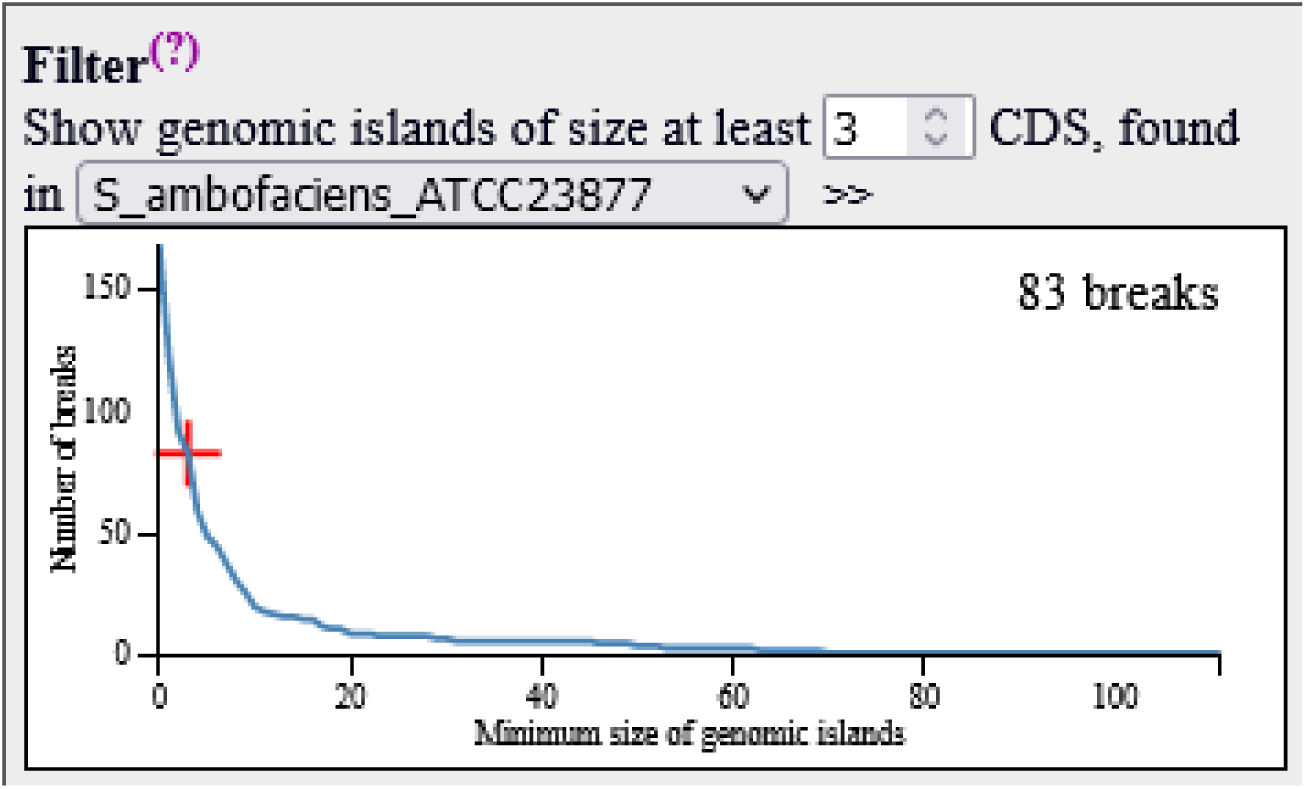
Number of genomic islands detected in *S. ambofaciens* ATCC23877 genome when compared to *S. coelicolor* A3(2) related to the minimal number of CDSs in the genomic islands.

### Global analysis of the genomic islands identified in the genome of *S. ambofaciens* ATCC 23877 when compared to the genome of *S. coelicolor* A(3)2

To allow the analysis of genomic islands, SYNTERUPTOR provides access to a comprehensive list of all identified genomic islands through a link on the dotplot page, leading to a detailed table. This table contains characteristics of the genomic islands determined by the program, including their location, their number of CDSs, the number of CDSs without ortholog in the second genome, the number of CDSs with paralogs in the considered genome, the presence of tRNA gene(s) within or near the genomic island, a brief description of the gene content, and the difference in GC ratio between the genomic island and the complete genomic sequence. Users have the option to download this table as a csv file for further analysis and to define weight coefficient to parameters (listed in Table S1) to rank genomic islands according to criteria of their choice for a customized and focused analysis.

In our specific comparison of the genomes of *S. ambofaciens* and *S. coelicolor*, with the minimum size of genomic islands set to three CDSs, SYNTERUPTOR detected 83 putative genomic islands in the *S. ambofaciens* genome. Of these, 46% contain five or fewer CDSs, primarily consisting of hypothetical proteins (40%), excreted proteins (11%), and membrane proteins (11%). The abundance of hypothetical proteins in these islands makes it difficult to hypothesize their putative functions.

Genomic islands comprising six to ten CDSs represent 32% of all islands, while those with 11 to 20 CDSs account for 11%. The remaining 11% consist of genomic islands containing 22 to 111 CDSs (Table 1). Among these larger islands, some contain known or putative mobile genetic elements (GI-1 and GI-3) or SMBGCs (GI-2 and GI-8). For instance, GI-3 contains the pSAM2 integrative conjugative element and the putative prophage xSAM1 (43), while the largest island (GI-1, 111 CDS) harbours the Samy prophage (44). GI-2 contains the spiramycin biosynthetic gene cluster (45), along with an SMGBC predicted to direct the biosynthesis of a lipopeptide related to the calcium-dependent antibiotic (CDA, (41)). GI-8 contains the SMBGC responsible for congocidine biosynthesis (46). This analysis confirms that SYNTERUPTOR is effective in finding SMBGCs when they are located within genomic islands. Moreover, it highlights the potential applications of SYNTERUPTOR beyond SMBGC discovery, such as the identification of mobile genetic elements and pathogenicity or symbiosis islands in other domains of biology.

**Table 1.** Genomic islands of more than 20 CDSs in the genome of *S. ambofaciens* ATCC23877 (genome comparison with *S. coelicolor* A(3)2)

| Name | Number of CDSs | Start (nt) | End (nt) | Genes within the genomic island | Putative content |
| --- | --- | --- | --- | --- | --- |
| GI-1 | 111 | 6 579 545 | 6 650 946 | SAM23877_6075 - SAM23877_6185 | putative prophage(s) |
| GI-2 | 69 | 5 969 372 | 6 128 438 | SAM23877_5592 - SAM23877_5660 | spiramycin/NRP SMBGC |
| GI-3 | 62 | 4 270 811 | 4 326 714 | SAM23877_4016 - SAM23877_4077 | pSAM2-xSAM1 |
| GI-4 | 52 | 6 965 819 | 7 022 659 | SAM23877_6462 - SAM23877_6513 | contains CRISPR system |
| GI-5 | 49 | 316 599 | 358 923 | SAM23877_0359 - SAM23877_0407 | unknown |
| GI-6 | 45 | 7 947 017 | 7 989 060 | SAM23877_7262 - SAM23877_7306 | unknown |
| GI-7 | 30 | 6 486 283 | 6 513 450 | SAM23877_5987 - SAM23877_6016 | unknown |
| GI-8 | 28 | 7 191 867 | 7 225 858 | SAM23877_6654 - SAM23877_6681 | Congocidine SMBGC |
| GI-9 | 22 | 4 171 706 | 4 190 932 | SAM23877_3918 - SAM23877_3939 | Sphydrofuran SMBGC |

### Detection of an uncharacterised SMBGC in GI-9

The analysis of genomic islands confirmed that SYNTERUPTOR has the ability to identify SMBGCs. However, the SMBGCs detected in this analysis were either previously known and characterised (spiramycin, congocidine) or easily detectable using similarity-based tools like antiSMASH. Among the remaining large genomic islands (>20 CDS), one specific island (GI-9) caught our attention. It harboured genes that encode enzymes potentially involved in the biosynthesis of a specialised metabolite. At the outset of our study, this particular genomic region was not detected by antiSMASH.

GI-9, identified as SYNTERUPTOR id 0762fd (Figure 2), consists of 22 CDSs (SAM23877_3918 to SAM23877_3939). Notably, it contains a gene (SAM23877_3931) that encodes a homolog of AvrD from *Pseudomonas syringae* and MmyD from *S. cœlicolor* A(3)2 (Table 2). These two butenolide synthases are known for their involvement in the biosynthesis of syringolides (plant defence elicitors) and the methylenomycin antibiotic, respectively (47, 48). Additionally, GI-9 includes other putative biosynthetic genes related to polyketide/fatty acid biosynthesis (SAM23877_3923, SAM23877_3929, SAM23877_3930, and SAM23877_3936). Moreover, there are genes possibly associated with transcriptional regulation (SAM23877_3919, SAM23877_3922, SAM23877_3934, SAM23877_3935 and SAM23877_37) or transport (SAM23877_3921). Considering the potential significance of GI-9 and its unreported detection by antiSMASH at the beginning of our study, we considered it as a promising candidate to assess whether SYNTERUPTOR could serve as a valuable tool for the discovery of novel SMBGCs.

**Table 2.** Gene content of GI-9 (Synteruptor id 0762fd)

| **Gene** | **Predicted function** |
| --- | --- |
| SAM23877_3918 | Polysaccharide deacetylase family protein |
| SAM23877_3919 | Helix-turn-helix transcriptional regulator |
| SAM23877_3920 | Uma2 family endonuclease |
| SAM23877_3921 | MFS transporter |
| SAM23877_3922 | Helix-turn-helix transcriptional regulator |
| SAM23877_3923 | Ketoacyl-ACP synthase III family protein |
| SAM23877_3924 | Aldo/keto reductase |
| SAM23877_3925 | Carboxymuconolactone decarboxylase family |
| SAM23877_3926 | FAD-dependent monooxygenase |
| SAM23877_3927 | Antibiotic biosynthesis monooxygenase |
| SAM23877_3928 | Histidine phosphatase family protein |
| SAM23877_3929 | Acyl carrier protein (ACP) |
| SAM23877_3930 | Beta-ketoacyl-ACP synthase III |
| SAM23877_3931 | AvrD family protein |
| SAM23877_3932 | Hotdog fold thioesterase |
| SAM23877_3933 | Aldo/keto reductase |
| SAM23877_3934 | LuxR C-terminal-related transcriptional regulator |
| SAM23877_3935 | Helix-turn-helix domain-containing protein |
| SAM23877_3936 | Beta-ketoacyl-[acyl-carrier-protein] synthase family protein |
| SAM23877_3937 | CopG family transcriptional regulator |
| SAM23877_3938 | PIN domain-containing protein |
| SAM23877_3939 | Hypothetical protein |

To investigate whether GI-9 contains a specialised metabolite biosynthetic gene cluster, we inactivated SAM23877_3931 in the OSC4 strain, yielding OSC416. OSC4 is a derivative of the wild type *S. ambofaciens* ATCC23877, lacking the pSAM2 integrated conjugative element (43), and unable to produce several known *S. ambofaciens* ATCC23877 antibacterial metabolites (spiramycins, congocidine, and kinamycins). Since the optimal culture conditions for expressing the GI-9 genes were unknown, we adopted an OSMAC approach (49) to explore the production of metabolite(s) potentially directed by the putative SMBGC within GI-9.

We cultivated both OSC4 and OSC416 strains in various media and subjected the culture supernatants to HPLC analyses. When the supernatant of OSC4 cultivated in R2YEm medium was analysed, two distinct metabolites, named M1 and M2 (retention times of 4.7 min and 12.5 min), were detected. In contrast, these metabolites were absent from the chromatogram of the supernatant of OSC416 cultivated under the same conditions (Figure 5). Similar results were observed when the strains were grown in R2 and R2YE media. This finding suggested that GI-9 indeed contained an uncharacterised SMBGC.

**Figure 5.**
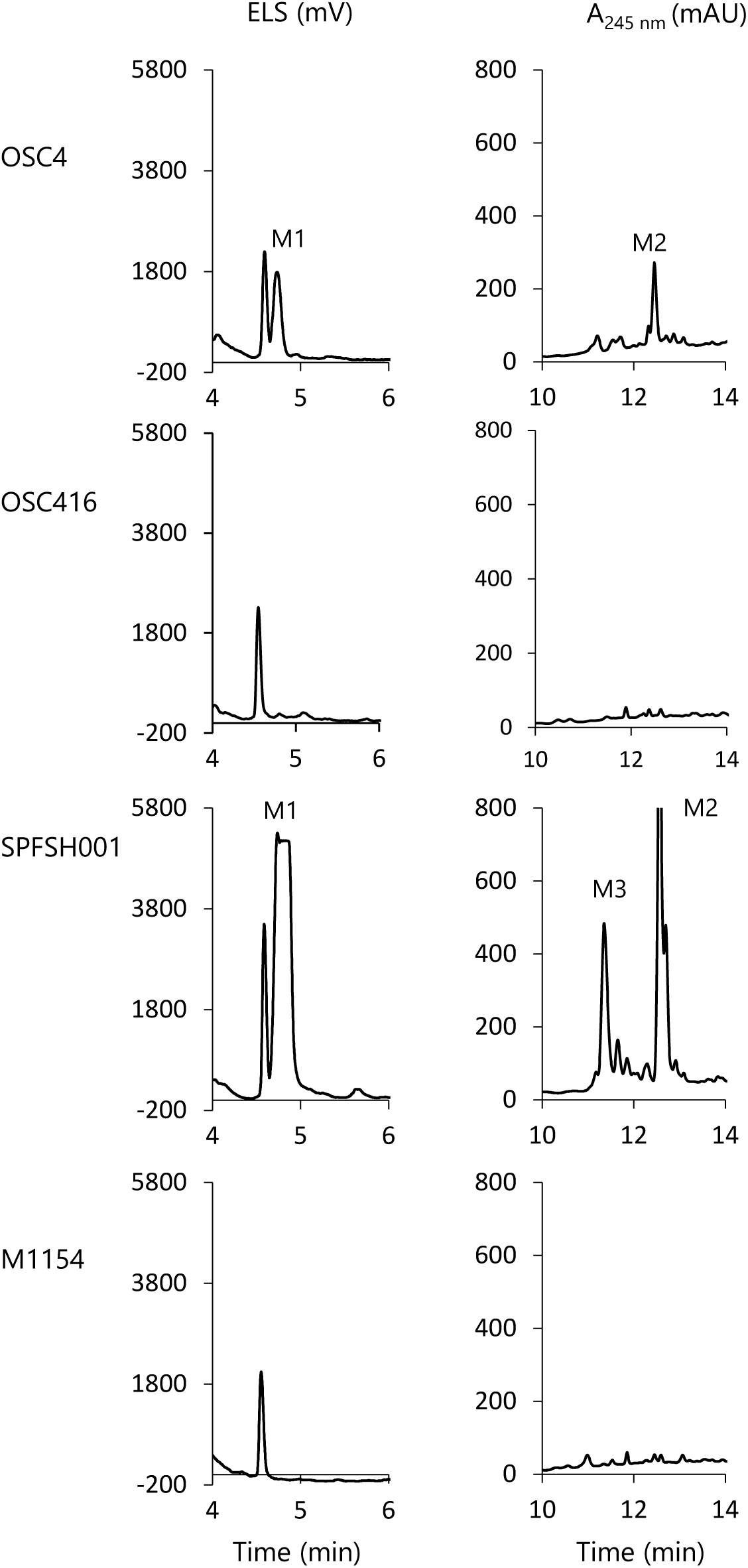
HPLC analysis of culture supernatants *of S. ambofaciens* ATCC 23877 OSC4 and OSC416 (SAM23877_3931 inactivated), *S. coelicolor* M1154 and *S. coelicolor* SPFSH001. First column, ELSD monitoring; second column UV monitoring at 245 nm.

To further validate the presence of a functional SMBGC in GI-9, we pursued the heterologous expression of GI-9. For this purpose, we constructed a cosmid library of the genomic DNA of *S. ambofaciens* ATCC23877 and screened it using an internal fragment of SAM23877_3931 as a probe. We isolated twelve cosmids from the library and screened them for the complete GI-9 region. By sequencing the ends of the inserts of two out of the four cosmids containing the entire GI-9, we determined their precise inserts. One of these cosmids, named pSLM003, was then introduced by intergeneric conjugation into the heterologous host *S. coelicolor* M1154, resulting in the strain SPFSH001. SPFSH001 and M1154 were cultivated in R2YEm for five days at 30°C and the culture supernatants were analysed by HPLC. Figure 5 shows that the two metabolites M1 and M2 are produced by SPFSH001 but not by the *S. coelicolor* M1154 host. Interestingly, a third metabolite, M3, was also observed in the culture supernatant of SPFSH001, metabolite that was absent in the supernatant of OSC16. These experimental results confirmed the presence of a SMBGC in GI-9.

### M1 purification and chemical characterization

We proceeded with the purification of the main compound, M1 (detected only by ELS). The supernatant from a 1-liter culture of SPFSH001 was evaporated to dryness under reduced pressure, yielding a residue that was then extracted with methanol. The methanol extract underwent flash chromatography, and fractions containing M1 were identified through TLC analysis and subsequently combined. To further purify M1, preparative RP-HPLC was performed under acidic conditions. After *in vacuo* concentration, we obtained 19 mg of M1 as a yellowish oil.

High resolution mass spectrometry (HR-ESI-MS *m/z* [M+H]^+^ 173.0828 (calcd for C_8_H_13_O_4_, 173.0814), Figure S1) and ^1^H and ^13^C NMR analyses (Figures S2 and S3, Table S4) identified M1 as 2-methyl-4-(1-glycerol)-furan (MGF, compound 2, Figure 6). This compound was previously isolated by Umezawa and colleagues in 1971 (50) as the hydrolysis product of sphydrofuran (compound 1, Figure 6), the structure of which was determined by Usui and colleagues (51). As M1 was purified under acidic conditions during preparative RP-HPLC, we investigated the possibility that M1 might actually be sphydrofuran that underwent hydrolysis during the purification process, leading to the formation of the purified and chemically characterised MGF. To address this, the fraction obtained after preparative RP-HPLC was neutralized with sodium hydroxide. Subsequent analysis by ESI-MS and ^1^H NMR (Figures S4 and S5) confirmed that M1 is indeed sphydrofuran. Although we could not obtain sufficient quantities of M2 and M3 for chemical characterization, the identification of M1 as sphydrofuran constitutes the proof of concept that SYNTERUPTOR can be used to detect genomic islands containing original SMBGCs.

**Figure 6.**
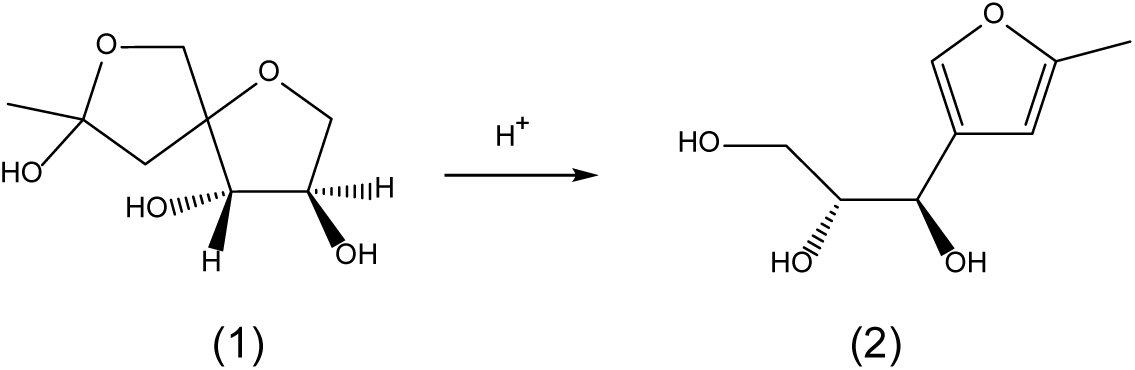
Scheme of the chemical degradation of sphydrofuran (1) into 2-methyl-4-(1-glycerol)-furan (2)

### Evaluating the Performance and Capabilities of SYNTERUPTOR

SYNTERUPTOR was specifically designed to compare the genome sequences of closely related species, accommodating the use of draft genome sequences as well. We wanted to assess how the program will perform with draft genome sequences and with relatively distantly related species. To achieve this, we created a SYNTERUPTOR database (https://bioi2.i2bc.paris-saclay.fr/synteruptor/summary.php?version=_mgnTDH0BR) containing the sequences of the chromosomes of 12 *Streptomyces* species in addition to the *S. ambofaciens* ATCC 23877 genome. These species exhibit average nucleotide identity with the genome of *S. ambofaciens* ATCC 23877 ranging from 98.98% (representing 87.3% of the genome sequence, *S. ambofaciens* DSM40697) to 75.59% (representing 33.1% of the genome sequence, *Streptomyces cattleya* DSM46488) (Table S5). While *S. ambofaciens* DSM 40697 clearly belongs to the same species as *S. ambofaciens* ATCC 23877, as determined by an ANI of ≥ 95%, ANI values for the remaining species are consistent with these species falling within the same bacterial genus (52). The number of contigs in genome sequences varies from one (complete genomes) to 660 (in the case of *Streptomyces* sp. alain-F2R5).

Supplementary Figure S6 illustrates the doptplots and GOC profiles generated by SYNTERUPTOR when comparing the genome sequences of these 12 *Streptomyces* species with the chromosome sequence of *S. ambofaciens* ATCC 23877. It shows that when working draft genome sequences, the total number of contigs should ideally be limited to a few dozen (Supplementary Figure S6-B, D and F), as exemplified by the genome sequence of *Streptomyces* sp. M1013 in our case (43 contigs, Supplementary Figure S6-D). Excessive contigs can hinder the accurate determination of syntenic regions, leading to potential errors in identifying synteny breaks.

Figure S6 and Table S5 also demonstrate that the number of genomic islands tends to increase with the phylogenetic distance between the compared species, as could be expected. Nonetheless, even when comparing strains of the same species, SYNTERUPTOR still identifies a notable number of genomic islands (e.g., 13 genomic islands consisting of 15 CDSs or more in the genome of *S. ambofaciens* ATCC 23877 when comparing the genomes of *S. ambofaciens* ATCC 23877 and DSM 40697). In this example, the nature of these islands may be suggestive of horizontal gene transfer events in most cases. Thus, two islands possess a putative integrase gene and a tRNA gene at one of their extremity, while five others contain one or several putative transposase genes. In addition, two islands are constituted by the xSAM1-pSAM2 AICE (53) and the Samy prophage (44). Interestingly, one of the genomic islands contains an SMBGC (iterative type I PKS).

The detected genomic islands are predominantly located in the central region of the chromosomes, where the GOC levels are relatively high. Some genomic islands can nonetheless be found towards the extremities of chromosomes, where GOC levels drop. However, these genomic islands should be approached with caution as they might arise from small synteny blocks (composed of two to three genes) within large regions where no significant synteny is otherwise observed (see Figure S7 as an example).

### Identification of hotspots of DNA integration

While SYNTERUPTOR was intended to aid in the identification of genomic islands with potential SMBGCs, it does not restrict itself solely to those regions. Instead, it identifies genomic islands in general, regardless of their content. Most of the time, these regions have been acquired through horizontal gene transfer. In a 2017 study, Oliveira *et al.* analysed the genomes of 80 bacterial species and observed that horizontally acquired DNA was concentrated in specific chromosomal regions known as “hotspots.” These hotspots represent approximately 1% of the chromosomal regions and are regions prone to acquiring genetic material through horizontal gene transfer (54). SYNTERUPTOR allows users to visualize these hotspots in their database. Indeed, the table describing the genomic islands includes a diversity factor. This factor represents the number of genomic islands found at this specific location in the other genomes included in the database. In addition, these hotspots can be visualized in each individual genomic island webpage, as a network and an alignment (Figure 7). In the network (break graph), the link between one species and another one indicates the existence of a synteny break at the exact same location in the chromosome. The alignment allows to visualize the content of each genomic island. In the given example, the genome of the strain *S. ambofaciens* DSM40697 does not contain any genomic island at the considered position, while the genomes of the strains *S. ambofaciens* ATCC23877, *S. coelicolor* A(3)2, *Streptomyces pactum* ACT12 and *Streptomyces* sp. FXJ7_023 each contain a distinct genomic island. These islands are located next to a tRNA-Ser encoding gene, suggesting that they may be the result of a horizontal gene transfer event.

**Figure 7.**
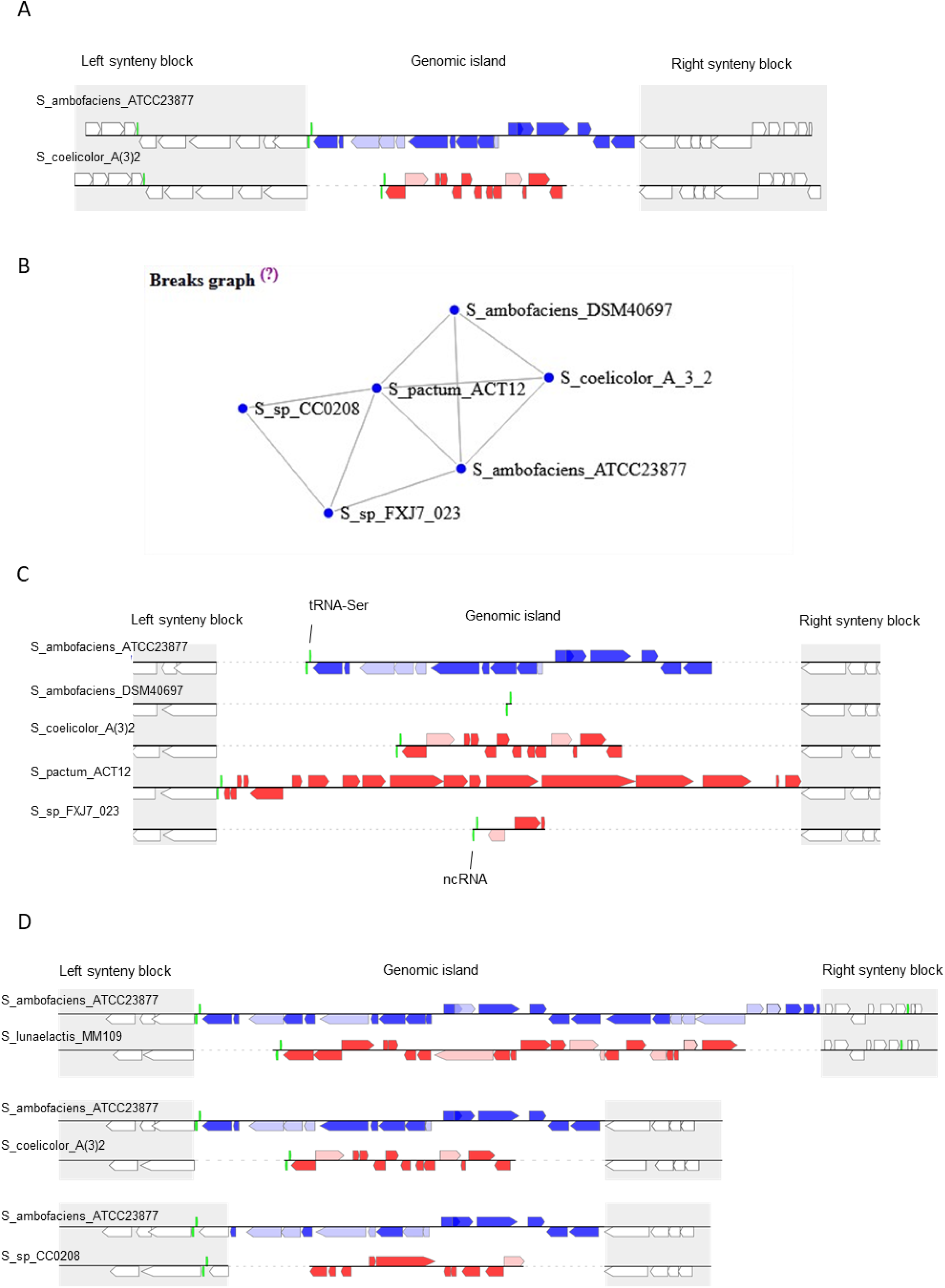
Example of a hotspot of integration detected by SYNTERUPTOR. A) Gene content and organization of the genomic islands in the synteny break 478e7e (#807); B) Break graph representing the existence of genomic islands (link between two organisms) located at the same position in the genomes of the database than the genomic islands observed in A; and C) Visualisation of the genomic islands located at the same position than the genomic islands observed in A. D) Illustration of overlapping breaks for the genomic island presented in A, showing that in *S. lunaelactis* MM109 and *Streptomyces* sp. CC0208 genomes, genomic islands are present in the same chromosomal region, but the boundaries of these islands are slightly different.

The break graph and the break alignment allow to visualise genomic islands that are defined by the exact same boundaries (Figure 7C). However, in some cases, a genomic island with slightly different boundaries may be present in same chromosomal region of a genome of the database. We named those genomic islands, which occur more frequently when comparing more distantly related species, “overlapping genomic islands” (identified in overlapping synteny breaks). They are presented next to the break graph and the break alignment and can be individually visualised. In the example presented in Figure 7, the genomes of *S. lunaelactis* MM109 and *Streptomyces* sp. CC0208 contain a genomic island in the same region as the ones located in *S. ambofaciens* ATCC23877 and *S. coelicolor* A(3)2 (synteny break #807), but with slightly different boundaries (Figure 7D).

### Complementarity with other bioinformatic tools

SYNTERUPTOR is most useful when comparing the genomes of relatively close species. To identify these species, it may be advantageous to make use of the autoMLST tool developed to perform rapid phylogenetic analyses (55).

Among the many tools developed to identify specialised metabolite gene clusters in bacterial genomes, antiSMASH is probably the most widely used, as it allows the detection of gene clusters directing the biosynthesis of a large diversity of specialised metabolites (17).

Therefore, we compared the genomic regions of *S. ambofaciens* ATCC23877 found with antiSMASH 6.0 (run in relaxed or loose mode) and with the genomic islands detected by SYNTERUPTOR (comparison with *S. coelicolor* A(3)2, Supplementary Tables 6-7-8). Among the 83 genomic islands detected by SYNTERUPTOR (Table 3), seven are located outside of the region of the chromosome highly syntenic with the chromosome of *S. coelicolor* A(3)2, estimated using both the dotplot and GOC profiles (roughly between SAM23877_0680 and SAM23877_7060). antiSMASH identified 27 (relaxed mode) and 49 (loose mode) putative gene clusters, with respectively 20 and 39 located in the region of the chromosome highly syntenic with the chromosome of *S. coelicolor* A(3)2.

**Table 3:**
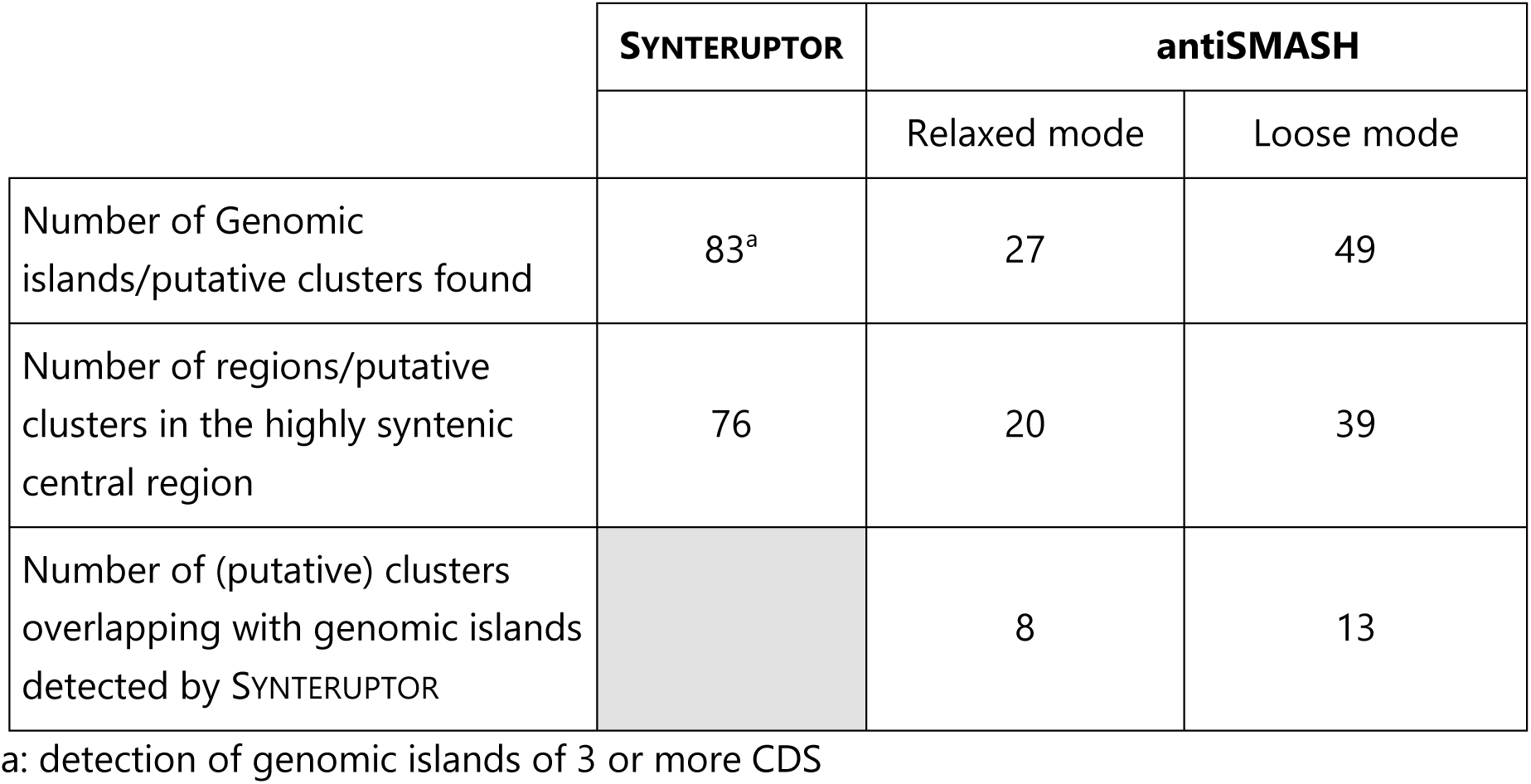
Genomic regions found with SYNTERUPTOR and with antiSMASH (Relaxed or loose mode)

The comparison of the genomic islands (SYNTERUPTOR) and the putative SMBGC regions (antiSMASH) showed that eight (relaxed mode) or 13 chromosomic regions (loose mode) are found by both programs (description in supplemental Table S8). Most of the time (for 11 of the 13 regions), the genomic islands identified by SYNTERUPTOR (and especially the small ones) are included within the SMBGCs regions proposed by antiSMASH. One of the reasons is that antiSMASH SMBGC regions can be quite large, as they may contain several SMBGCs, as it is the case for Regions 2 (2-methylisoborneol and antimycin SMBGCs) or 31 (spiramycin and a NRPS SMBGCs). In these cases, SYNTERUPTOR can be useful to better define the limits of the SMBGCs found by antiSMASH. For example, Region 39 identified by antiSMASH (41 CDSs) contains the congocidine SMBGC (22 CDSs). GI #838 (included in Region 39) allows to narrow down the region containing this cluster to 28 genes. The help that SYNTERUPTOR can provide to identify individual SMBGCs in antiSMASH regions is even clearer when looking at Region 20. This region contains 99 CDSs, and antiSMASH predicts four candidate gene clusters (two single ones, one interleaved and one neighbouring, Figure S8). GI #819 allows to delimit the interleaved one to 22 CDSs. This genomic island contains the sphydrofuran SMBGC (constituted of 12 CDS, SAM23877_3924 – SAM23877_3936, our unpublished results).

Among the SMBGCs not detected by SYNTERUPTOR are found most of the gene clusters constituting the core specialised metabolism of *Streptomyces* species, such as the desferrioxamine, albaflavenone, ectoine or isorenieratene SMBGCs. These clusters, conserved among *Streptomyces*, are found within regions of the chromosome of *S. ambofaciens* syntenic with the chromosome of *S. coelicolor* A(3)2 (Figure 8). These well-known SMBGCs would probably not be found either using SYNTERUPTOR with other *Streptomyces* genomes, but do not present a major interest when mining genomes for SMBGCs. More interesting are the strain-specific SMBGCs, which, for *S. ambofaciens* ATCC 23877, includes spiramycin, congocidine, kinamycin, antimycin, stambomycin and sphydrofuran SMBGCs. These SMBGCs could potentially be detected by SYNTERUPTOR as they are absent from the genome of *S. coelicolor* A(3)2. Indeed, three of them (spiramycins, congocidine and sphydrofuran) are detected. However, the kinamycin, antimycin and stambomycin SMBGCs fail to be detected by SYNTERUPTOR. These clusters are located at the extremities of the chromosome of *S. ambofaciens* ATCC23877, in regions where the synteny with the chromosome of *S. coelicolor* A(3)2 is too low for their detection (Figure 8).

**Figure 8.**
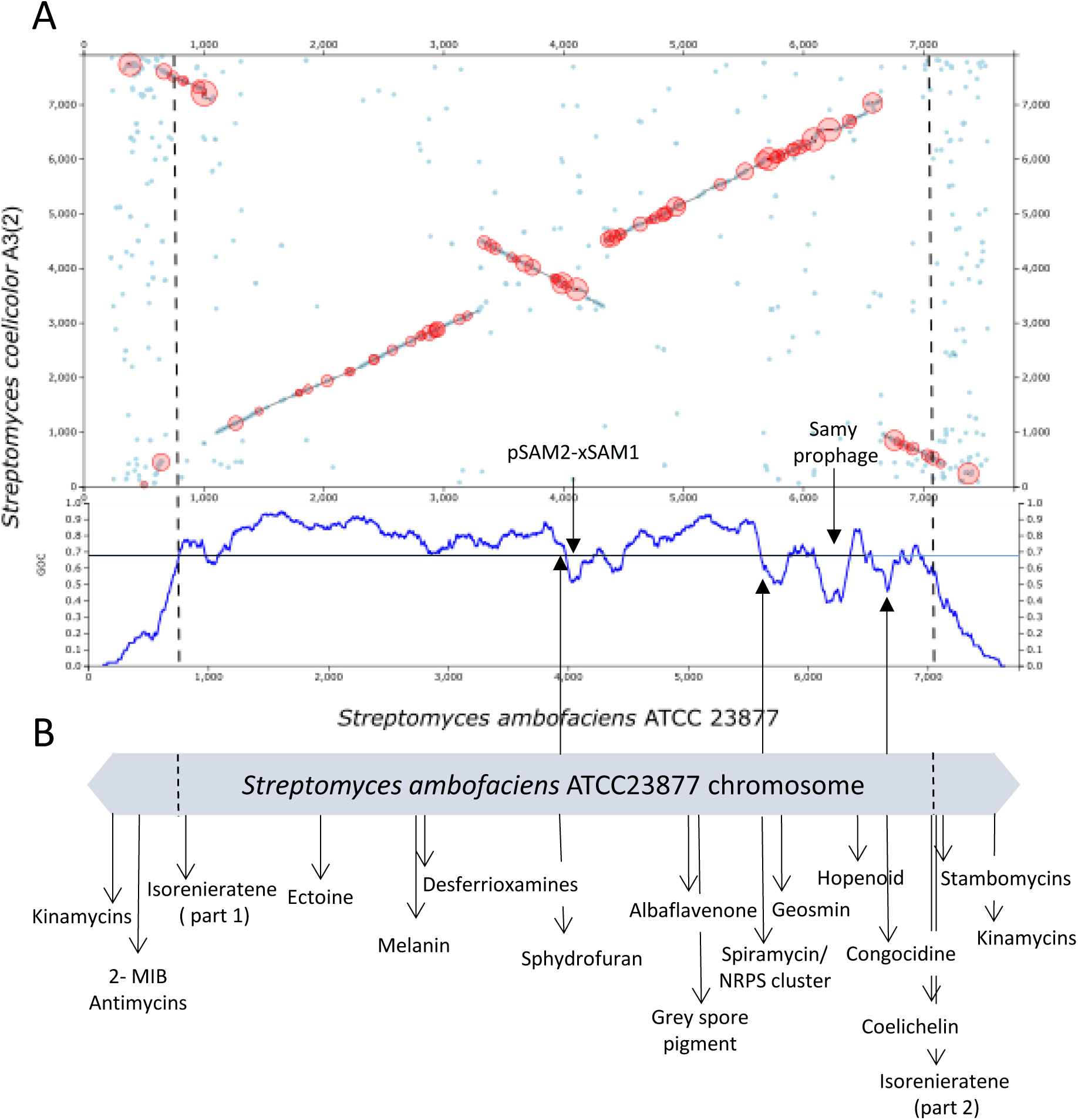
Localisation of *S. ambofaciens* ATCC 23877 genomic islands and SMBGCs; A. Screenshot of SYNTERUPTOR showing the dotplot and GOC profile when comparing *S. ambofaciens* ATCC23877 and *S. coelicolor* A(3)2 chromosomes and looking for genomic islands of three or more CDS in *S. ambofaciens*. Dotted lines indicate the approximate limits of the synteny region with the chromosome of *S. coelicolor* A(3)2.; B. Schematic representation of the *S. ambofaciens* ATCC23877 chromosome with the location of the known SMBGCs and of the newly identified one, sphydrofuran.

Mining bacterial genomes for SMBGCs is not always a straightforward task, particularly when dealing with biosynthetic enzymes belonging to underexplored or poorly families, as it was the case until recent years, for example, for enzymes catalysing the formation of N-N bonds. With SYNTERUPTOR, we present the specialised metabolism community with a bioinformatic tool that could be useful in the exploration of bacterial genomes for SMBGCs that may go unnoticed by conventional tools like antiSMASH. In this context, we demonstrate the utility of SYNTERUPTOR by identifying a genomic island housing the sphydrofuran SMBGC. As antiSMASH continues to enhance its cluster detection capabilities, SYNTERUPTOR could remain a valuable resource, aiding in the delineation of individual SMBGCs within antiSMASH regions that may encompass multiple clusters and in refining the boundaries of these SMBGCs. Finally, the capability of SYNTERUPTOR to identify genomic islands, regardless of their contents, extends its applicability to other biological contexts, including the investigation of various types of functional genomic islands and the examination of events related to horizontal gene transfer.

## DATA AVAILABILITY

The SYNTERUPTOR program can be freely accessed at https://bioi2.i2bc.paris-saclay.fr/synteruptor. The SYNTERUPTOR source code is available in Zenodo [10.5281/zenodo.10424133], at [DOI 10.5281/zenodo.10424123].

## SUPPLEMENTARY DATA

Supplementary Data are available online.

## ACKNOWLEDGEMENTS

We acknowledge the Integrative Bio Informatics (BioI2) platform of the I2BC and especially Chloé Quignot for hosting and maintaining the SYNTERUPTOR program. During the preparation of this manuscript, the authors used the chatGPT service to rephrase some sentences and improve the overall quality of the English language.

