## Supplementary methods, Tables and Figures for "SYNTERUPTOR: mining genomic islands for non-classical specialised metabolite gene clusters"

### SUPPLEMENTARY MATERIAL

**SUPPLEMENTARY METHODS**

Construction of the OSC4 mutant 3

Construction of OSC416 mutant 4

**SUPPLEMENTARY TABLES**

Supplementary Table S1: List of parameters used for ranking genomic islands 5

Supplementary Table S2: Strains, plasmids and cosmids used in this study 6

Supplementary Table S3: Oligonucleotides used in this study 7

Supplementary Table S4: NMR Spectroscopic data of 2-methyl-4-(1-glycerol)-furan, 8

Supplementary Table S5: Characteristics of the genome sequences of *Streptomyces* species used to construct the SYNTERUPTOR database 9

**SUPPLEMENTARY FIGURES**

Supplementary Figure S1: HR-ESI-MS-analysis of purified metabolite M1 under acidic conditions 10

Supplementary Figure S2: 1H-NMR spectrum (500 MHz, MeOD) of purified metabolite M1 under acidic conditions 11

Supplementary Figure S3: 13C-NMR spectrum (125 MHz, MeOD) of purified metabolite M1 under acidic conditions 12

Supplementary Figure S4: MS-ESI (negative mode) analysis of the neutralized HPLC fraction 13

Supplementary Figure S5: 1H NMR spectra of the neutralized HLC fraction 14

Supplementary Figure S6: GOC profiles and doptplots generated by SYNTERUPTOR when comparing the genome sequences of 12 *Streptomyces* species with the genome sequences of *S. ambofaciens* ATCC 23877 15

Supplementary Figure S7: Example of a genomic island (cd1336) obtained by SYNTERUPTOR, delimited by synteny regions constituted of two or three CDS in an otherwise large non-syntenic region. 21

Supplementary Figure S8: Visualisation of Region 20 of *S. ambofaciens* ATCC 23877 genome predicted to contain one or several SMBGCs. 22

**REFERENCES** **23**

#### Supplementary methods

##### Construction of the OSC4 mutant

The OSC4 mutant was constructed to abolish the production of metabolites with known antibacterial activities, i.e. spiramycins, congocidine and kinamycins by *S. ambofaciens* ATCC23877. First, the *alpID* locus (5 genes: *alpI*, *alpA*, *alpB*, *alpC* and *alpD*) from the kinamycin biosynthetic gene cluster was deleted by PCR targeted mutagenesis (1) as previously described (2). Briefly, the cosmid F6Δ*alpID*::*aac(3)*IV-oriT (*alpID* locus replaced by the *aac(3)*IV-oriT cassette (2) was introduced into *E. coli* ET12567/pUZ8002 and then transferred into *S. ambofaciens* OSC2 (3) by intergenic conjugation on SFM $MgCl_2$ (10 mM). Exconjugants were selected by overlaying the plates with apramycin (final concentration 50 mg/ml) and nalidixic acid (final concentration 25 mg/ml) to eliminate *E. coli* cells. The clones in which the locus replacement has occurred (double-crossover exconjugants) were selected on their phenotype ([$Kan^S$, $Apra^R$]). As the kinamycin cluster is located in the terminal inverted repeats of the chromosome, the *alpID* locus is then present in two copies on the chromosome. To ensure that the two loci were deleted and replaced by the cassette, the exconjugants [$Kan^S$, $Apra^R$] were tested for their abilities to produce a diffusible orange pigment (a pigment always associated to the production of kinamycin, (2)). The mutants unable to produce the pigment were analysed by Southern blot analysis using F6 DNA as probe and by PCR with the pair of primers CK1/CK2 as described (2) to confirm the gene replacement in the two chromosomal arms. The *aac(3)*IV-oriT cassette which is flanked by the FRT sites (1, 2) was then removed in the ΔΔ*alpID*::*aac(3)*IV-oriT strain by site specific recombination using the pUWLFLP plasmid which encodes a Flp recombinase as described in (4). The genotype of the ΔΔ*alpID*::scar clones was checked by PCR using CK1 and CK2 as primers. One of the clones was named H1 and used for further deletions.

To abolish the production of spiramycins in the H1 mutant, the pathway-specific activator *srm*40 was replaced by an apramycin resistance cassette. For this purpose, we used the cosmid pSPM107 (5). This cosmid contains part of the spiramycin gene cluster with the *srm*40 gene replaced by an apramycin resistance excisable cassette. The cosmid was introduced in *S. ambofaciens* H1 by conjugation using *E. coli* S17.1/pSPM107. Exconjugants were selected for apramycin resistance. Double recombinants were obtained by screening the apramycin resistant clones for puromycin sensitivity. The resulting clones were verified by PCR using primers KF64 and KF65 and named SPM700. The loss of spiramycin production was verified by bioassays as previously described (6).

To abolish congocidine production, the DNA fragment encompassing *cgc*22 to *cgc*18 was replaced by a hygromycin resistance excisable cassette. A 1.1 kb fragment internal to *cgc*22 was amplified using the CG3+f and CG3+r oligonucleotides and pCGC001 as template and cloned into pGEM-T Easy. The resulting plasmid was named pCGC507. Similarly, a 1.5 kb fragment overlapping *cgc*18 and *cgc*19 was amplified using the CG18_SNf and CG18_SNr oligonucleotides and cloned into pGEM-T Easy. The resulting plasmid was named pCGC509. The 1.1kb EcoRI/XbaI DNA fragment of pCGC507 and the 1.5 kb XbaI/HindIII fragment of pCGC509 were cloned into the EcoRI/*Hind*III digested pOJ260, yielding pCGC515. The hygromycin resistance cassette *att3Ωhyg* was obtained from pOSV709 (3) by EcoRV digestion and cloned into the XbaI/Klenow digested pCGC515, yielding pCGC516. The replacement of the *cgc* gene cluster by the *att3Ωhyg* cassette was obtained by introducing pCGC516 into *S. ambofaciens* H1 by conjugation using E. coli S17.1 and by selection of hygromycin resistant and apramycin sensitive clones. The resulting clones were verified by PCR using primer pairs (CGC18_SNf; R-aac2f), and (CG3+r; L-aac2r) and named CGCA008. The abolition of congocidine production was verified by bioassays on solid HT medium using *Bacillus mycoides* and *E. coli* as indicator strains.

A mutant unable to produce kinamycin, congocidine and spiramycins was obtained by mating the SPM700 and CGCA008 strains. Mutant clones were selected for resistance to apramycin and hygromycin. They were verified by PCR using the primer pairs (SN64f; SN65r), (CGC18_SNf; R-aac2f) and (CG3+r; L-aac2r) and the strain named OSC3. The *att3-aac* and the *att3Ωhyg* cassettes were excised using established methods (3) and the resulting clone was named OSC4.

**Construction of OSC416 mutant**

A 0.6 kb internal fragment of SAM23877_3931 was amplified by PCR from *S. ambofaciens* ATCC 23877 genomic DNA using the oligonucleotides mmyD-F and mmyD-R and cloned into the pSC-A vector (Agilent Technologies). The resulting plasmid was verified by restriction digestion and sequencing and named pAG001. The 0.6 kb EcoRI DNA fragment of pAG001 was next cloned into EcoRI-digested pOJ260 yielding pmmyD. This plasmid was introduced into *S. ambofaciens* OSC4 by conjugation with *E. coli* S17.1. Conjugants were selecting for apramycin resistance. The resulting SAM23877_3931 insertion mutant was verified by PCR using the primer pairs (up mmyD; lacZ-1) and (down mmyD; lacI-2) and named OSC416.

**Supplementary Table S1**: List of parameters used for ranking genomic islands

| Field | Description |
|---|---|
| tRNA genes adjacent to the genomic island | Presence of tRNA gene(s) at an extremity of the genomic island in one or both genomes |
| Diversity | Number of genomic islands found at the same location in other genomes of the database |
| Number of CDSs in the genomic island of genome 1 | Total number of CDSs found in the genomic island of genome 1 |
| Number of CDSs in the genomic island of genome 2 | Total number of CDSs found in the genomic island of genome 2 |
| Number of CDSs in the genomic island of genome 1 without orthologs in genome 2 | Number of CDSs in the genomic island in genome 1 without any orthologs in the genome 2 |
| Number of CDSs in the genomic island of genome 2 without orthologs in genome 1 | Number of CDSs in the genomic island in genome 2 without any orthologs in the genome 1 |
| Number of CDSs in the genomic island of genome 1 with at least one paralog | Number of CDSs in genome 1 with at least one paralog in the genome |
| Number of CDSs in the genomic island of genome 2 with at least one paralog | Number of CDSs in genome 2 with at least one paralog in the genome |
| Diff GC ratio in genome 1 | Difference of the GC ratio of the genes of this breaks with the GC ratio of the genome 1 |
| Diff GC ratio in genome 2 | Difference of the GC ratio of the genes of this breaks with the GC ratio of the genome 2 |

**Supplementary Table S2**: Strains, plasmids and cosmids used in this study

| Strains | Description | Source/Réference |
|---|---|---|
| ***Escherichia coli* strains** | | |
| S17.1 | Host strain for conjugation from *E. coli* to *Streptomyces* | Simon et al., 1983 (7) |
| ET12567/pUZ8002 | Nonmethylating strain with mobilization plasmid for conjugation with *Streptomyces* | Gust et al., 2004 (8) |
| ***Streptomyces ambofaciens*** | | |
| ATCC 23877 | Wild type strain | ATCC |
| OSC2 | ATCC23877 strain devoid of pSAM2 | Raynal et al., 2006 (3) |
| H1 | OSC2 in which the *alpID* loci involved in the kinamycin production are deleted | This study |
| SPM700 | H1 with *srm*40 replaced by the att3-*aac* cassette. | This study |
| CGCA008 | H1 with the *cgc* cluster replaced by the *att3Ωhyg* cassette | This study |
| OSC3 | H1 with *srm*40 replaced by the *att3-aac* cassette and the *cgc* cluster replaced by the *att3Ωhyg* cassette | This study |
| OSC4 | OSC3 strain in which the resistance cassettes have been removed. Mutant defective in the production of spiramycins/congocidine/ kinamycins | This study |
| OSC416 | Derived from OSC4, with SAM23877_3931 disrupted | This study |
| ***Streptomyces coelicolor*** | | |
| *Streptomyces coelicolor* M1154 | *S. coelicolor* M145 : Δact Δred Δcpk Δcda rpoB(C1298T) *rpsL*(A262G) | Gomez-Escribano & Bibb, 2011 (9) |
| SPFSH001 | *S. coelicolor* M1154 bearing pSLM003 | This study |
| **Plasmids/cosmids** | **Description** | **Source/Reference** |
| pSPM107 | Cosmid with the puromycin resistance gene and carrying part of the *srm* gene cluster including *srm40::att3-aac* | Karray et al., 2010 (5) |
| F6ΔalpID::aac(3)IV-oriT | F6 cosmid in which the *alpID* locus replaced by the *aac*(3)IV-*oriT* cassette | Pang et al., 2004 (2) |
| pOJ260 | Suicide vector for gene disruption in *Streptomyces* | Bierman et al., 1992 (10) |
| pCG001 | pBeloBAC11 containing a 43.4 kb *S. ambofaciens* DNA fragment carrying the entire congocidine gene cluster | Juguet et al., 2009 (11) |
| pCGC507 | 1.1 kb fragment internal to *cgc22* cloned into pGEM-T Easy | This study |
| pCGC509 | 1.5 kb fragment overlapping *cgc18* and *cgc19* cloned into pGEM-T Easy | This study |
| pCGC515 | pOJ260 containing the 1.1 kb *Eco*RI/*Xba*I DNA fragment of pCGC507 and the 1.5 kb *Xba*I/*Hind*III fragment of pCGC509 | This study |
| pCGC516 | *att3Ωhyg* cassette cloned in the XbaI/Klenow digested pCGC515 | This study |
| pOSV709 | pBCSK containing the *att3Ωhyg* cassette | Raynal et al., 2006 (3) |
| pSC-A | Cloning vector | Agilent Technologies |

| | | |
|---|---|---|
| pAG001 | pSC-A containing a 0.6 kb DNA fragment internal to SAM23877_3931 | This study |
| pmmyD | pOJ206 containing a 0.6 kb DNA fragment internal to SAM23877_3931 | This study |
| pWED4 | Cosmid derived from pWED1 with *hyg-oriT* and the ΦC31 integrative system | Vingadassalon et al., 2015 (12) |
| pSLM003 | pWED4 cosmid containing the complete genomic island GI-9 (id 0762fd) | This study |
| pSLM010 | pWED4 cosmid containing the complete genomic island GI-9 (id 0762fd) | This study |

**Supplementary Table S3**: Oligonucleotides used in this study

| Primer | Sequence 5'-3' | Description |
|---|---|---|
| KF64 | GGGACCACACCCGGCAGGACGACGG | Verification of the deletion of *srm40* in SPM700 |
| KF65 | ACGCACGCTTCGACCCGCCGGCGCG | Verification of the deletion of *srm40* in SPM700 |
| CG3+f | ATGCTCTAGAACGAGGAAGAGGACCCGCTCGACG | Amplification of an 1.1 kb fragment internal to *cgc*22, *Xba*I site underlined |
| CG3+r | TCCCGAATTCGGCCGGACCTTGCCCATGGAGTTC | Amplification of an 1.1 kb fragment internal to *cgc*22, *Eco*RI site underlined |
| CG18_SNf | ATCCAAGCTTTCGTCGGGTTGTCCGCATGGTTC | Amplification of a 1.5 kb fragment overlapping *cgc18* and *cgc19*, *Hind*III site underlined |
| CG18_SNr | TCAGTCTAGACATACGCCAGCGCACGGTGTTC | Amplification of a 1.5 kb fragment overlapping *cgc18* and *cgc19*, *Xba*I site underlined |
| R-aac2f | GGCCGTGACTGAGGAGGTCTAC | Verification of the CGCA008 mutant |
| L-aac2r | CCTCTCAGTCATGCGGGCAAC | Verification of the CGCA008 mutant |
| SN64f | GATGACGAGGCCCCCACGAGGACTCCTATG | Verification of the OSC3 mutant |
| SN65r | CAGGACGTGAGGTCGGTCAGCCACAGGTCG | Verification of the OSC3 mutant |
| mmyD-F | GCCGACTGGTCGAAGAAG | Inactivation of SAM23877_3931 |
| mmyD-R | GTCGTCGAGGTCGTAGAG | Inactivation of SAM23877_3931 |
| up mmyD | CCAGGCGTCGGACGTGTCCA | Verification of the OSC416 mutant |
| down mmyD | CGCCAATGCCGTCCTCGCGG | Verification of the OSC416 mutant |
| lacZ-1 | GGATGTGCTGCAAGGCGATT | Verification of the OSC416 mutant |
| lacI-2 | GCTTCCGGCTCGTATGTTGT | Verification of the OSC416 mutant |
| SL24 | GTCCGCATCGATCCCTAC | Amplification of an internal fragment of SAM23877_3940 |
| SL25 | ACCTGGGTCCGATGGAAG | Amplification of an internal fragment of SAM23877_3940 |
| SL26 | ATGAACGCCATCGACTCC | Amplification of an internal fragment of SAM23877_3916 |
| SL27 | GCGAACTCGACGTACTTG | Amplification of an internal fragment of SAM23877_3916 |

**Supplementary Table S4**: NMR Spectroscopic data[a] of 2-methyl-4-(1-glycerol)-furan

| | Sphydrofuran | |
|---|---|---|
| No. | $\delta_C$ (ppm) | $\delta_H$ (mult., *J* in Hertz) |
| 1 | 153.7 | - |
| 2 | 139.5 | 7.30 (1H, s) |
| 3 | 128.6 | - |
| 4 | 106.1 | 6.05 (1H, s) |
| 5 | 76.4 | 3.67 (1H, m) |
| 6 | 68.3 | 4.53 (1H, d, *5.96*) |
| 7 | 64.2 | 3.58 (1H, dd, *4.2*, *11.2*); 4.43 (1H, dd, *6.2*, *11.2*) |
| 8 | 13.4 | 2.24 (3H, s) |

[a] $^1$H chemical shifts were recorded at 500 MHz and $^{13}$C chemical shifts at 125MHz in MeOD

**Supplementary Table S5**: Characteristics of the genome sequences of *Streptomyces* species used to construct the SYNTERUPTOR database

| | Reference | ANIb and aligned percentage[a] | % synteny[b] | Number of contigs | Number of GIs of 15 or more CDS[c] |
|---|---|---|---|---|---|
| *Streptomyces ambofaciens* ATCC 23877 | NZ_CP012382.1 | | | **1** | |
| *Streptomyces ambofaciens* DSM 40697 | NZ_CP012949.1 | 98.98 [87.27] | 100 | 1 | 18 |
| *Streptomyces* sp. alain-F2R5 | NZ_NQOS01000100.1 | 88.34 [65.68] | 94 | 660 | 12 |
| *Streptomyces pactum* ACT12 | NZ_CP019724.1 | 87.98 [63.00] | 96 | 1 | 31 |
| *Streptomyces* sp. M1013 | NZ_MQUH01000001.1 | 87.11 [62.38] | 94 | 43 | 33 |
| *Streptomyces coelicolor* A3(2) | NC_003888.3 | 86.77 [57.28] | 94 | 1 | 35 |
| *Streptomyces* sp. FXJ7.023 | NZ_APIV01000001.1 | 85.54 [55.62] | 92 | 83 | 36 |
| *S. fungicidicus* TXX_3120 | NZ_CP023407.1 | 82.74 [53.72] | 91 | 1 | 35 |
| S. sp. CC0208 | NZ_CP031969.1 | 81.38 [50.66] | 89 | 1 | 57 |
| *S. lunaelactis* MM109 | NZ_CP026304.1 | 77.24[41.07] | 82 | 1 | 69 |
| *S. platensis* ATCC 23948 | NZ_CP023691.1 | 76.16 [38.12] | 78 | 1 | 56 |
| *S. bingchenggensis* BCW-1 | NC_016582.1 | 76.11 [41.46] | 75 | 1 | 88 |
| *Streptomyces cattleya* DSM 46488 | NC_017586.1 | 75.59 [33.10] | 79 | 1 | 53 |

a: ANIb = ANI blast; values given in comparison to the genome sequence of *S. ambofaciens* ATCC 23877

b: number of orthologs in synteny blocks / number of orthologs between two genomes; % given in comparison to the genome of *S. ambofaciens* ATCC 23877

c: in either genome

**Supplementary Figure S1**: HR-ESI-MS-analysis of purified metabolite M1 under acidic conditions

**Elemental Composition Report** **Page 1**

**Single Mass Analysis**
Tolerance = 30.0 PPM / DBE: min = -1.5, max = 100.0
Element prediction: Off
Number of isotope peaks used for i-FIT = 9

Monoisotopic Mass, Even Electron Ions
14 formula(e) evaluated with 1 results within limits (up to 50 closest results for each mass)
Elements Used:
C: 0-10 H: 0-20 O: 0-10
OUAZZANI_glegoff18-1 20 (0.532) Cm (20:24) 1: TOF MS ES+
1.37e+004

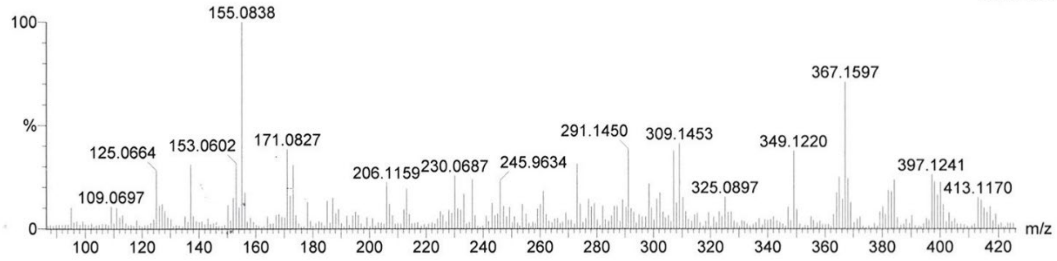


| | | | | | | | |
|---|---|---|---|---|---|---|---|
| Minimum: | | | | -1.5 | | | |
| Maximum: | | 5.0 | 30.0 | 100.0 | | | |
| Mass | Calc. Mass | mDa | PPM | DBE | i-FIT | i-FIT (Norm) | Formula |
| 173.0828 | 173.0814 | 1.4 | 8.1 | 2.5 | 116.6 | 0.0 | C8 H13 O4 |

**Supplementary Figure S2:** $^1$H-NMR spectrum (500 MHz, MeOD) of purified metabolite M1 under acidic conditions

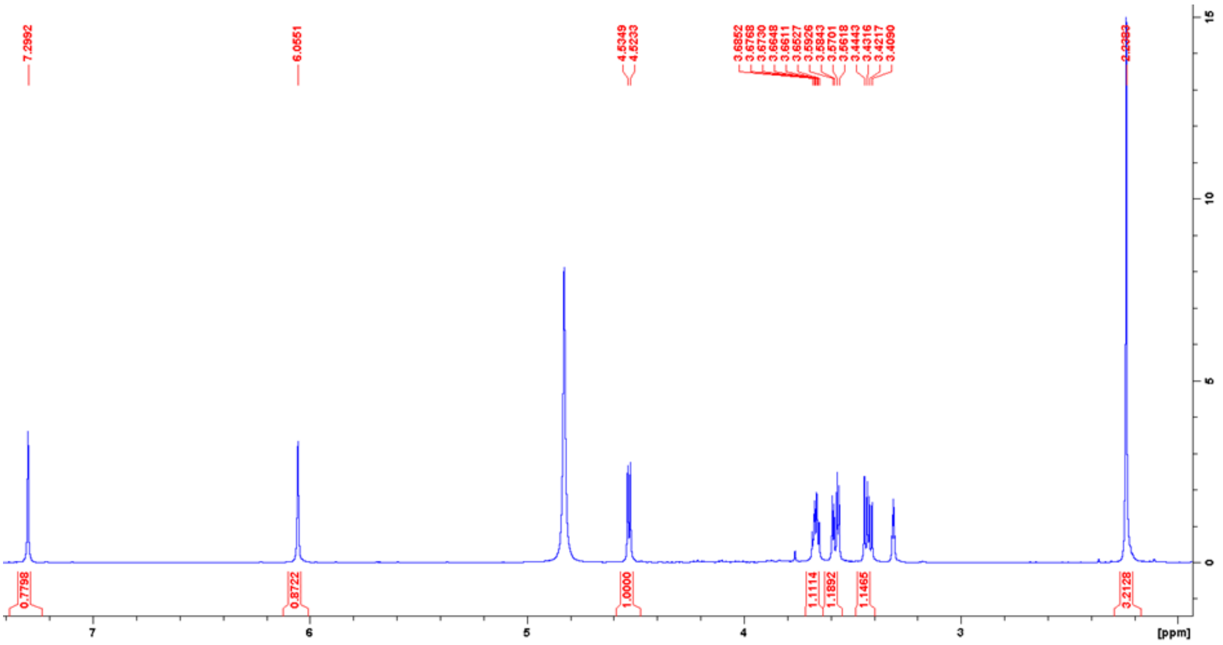

**Supplementary Figure S3:** $^{13}C$-NMR spectrum (125 MHz, MeOD) of purified metabolite M1 under acidic conditions

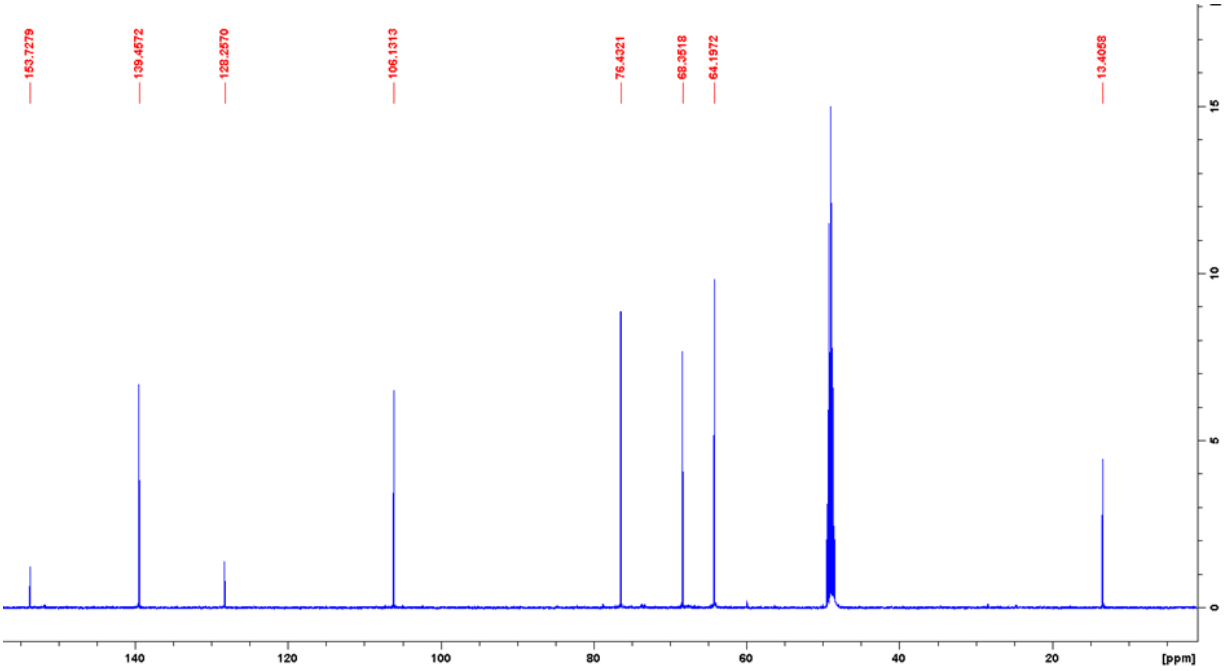

**Supplementary Figure S4:** MS-ESI (negative mode) analysis of the neutralized HPLC fraction

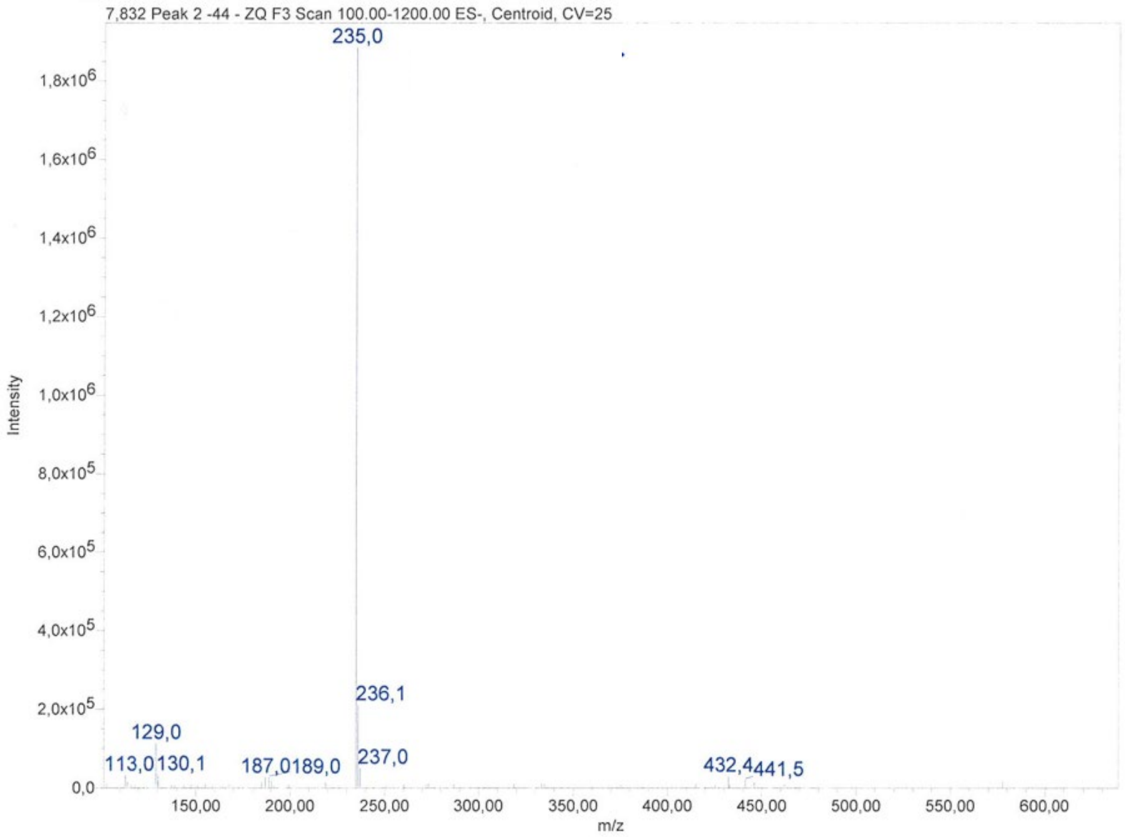


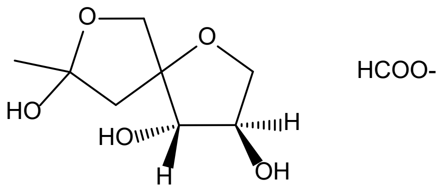

**Supplementary Figure S5**: 1H NMR spectra of the neutralized HLC fraction

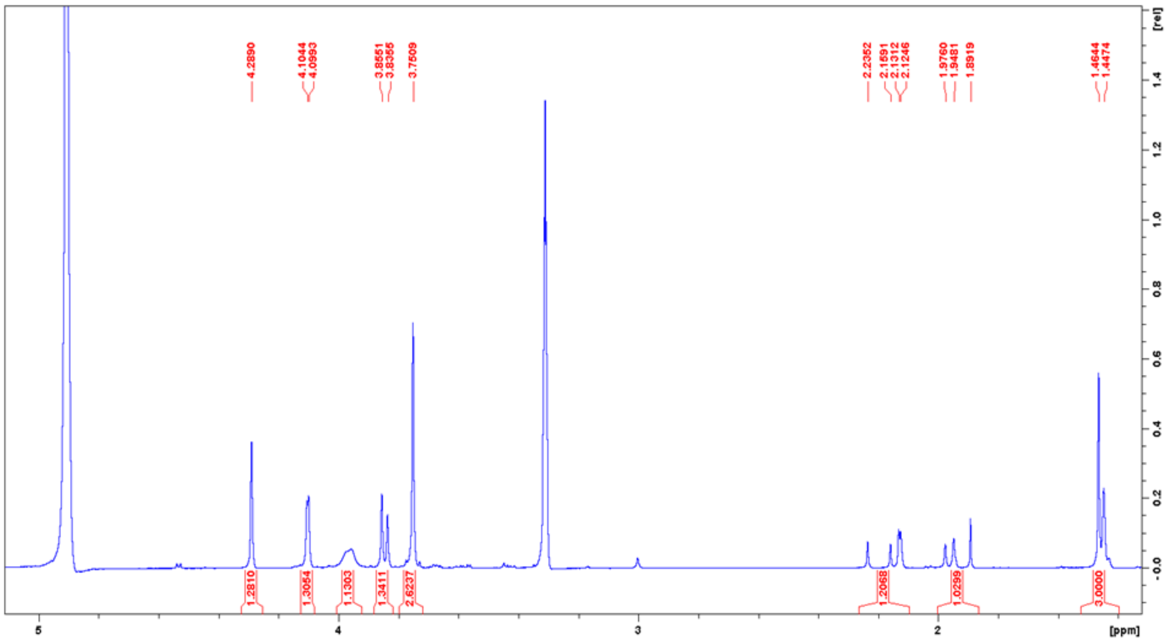

**Supplementary Figure S6**: GOC profiles and dotplots generated by SYNTERUPTOR when comparing the genome sequences of 12 *Streptomyces* species with the genome sequence of *S. ambofaciens* ATCC 23877. Only genomic islands of at least 15 CDSs found in at least one genome are shown. Grey horizontal lines in dotplots in B, D and F indicates the limits of the contigs.

A: *Streptomyces ambofaciens* DSM 40697
B: *Streptomyces* sp. alain-F2R5
C: *Streptomyces pactum* ACT12
D: *Streptomyces* sp. M1013
E: *Streptomyces coelicolor* A3(2)
F: *Streptomyces* sp. FXJ7.023
G: *Streptomyces fungicidicus* TXX_3120
H: *Streptomyces* sp. CC0208
I: *Streptomyces lunaelactis* MM109
J: *Streptomyces platensis* ATCC 23948
K: *Streptomyces bingchenggensis* BCW-1
L: *Streptomyces cattleya* DSM 46488

A

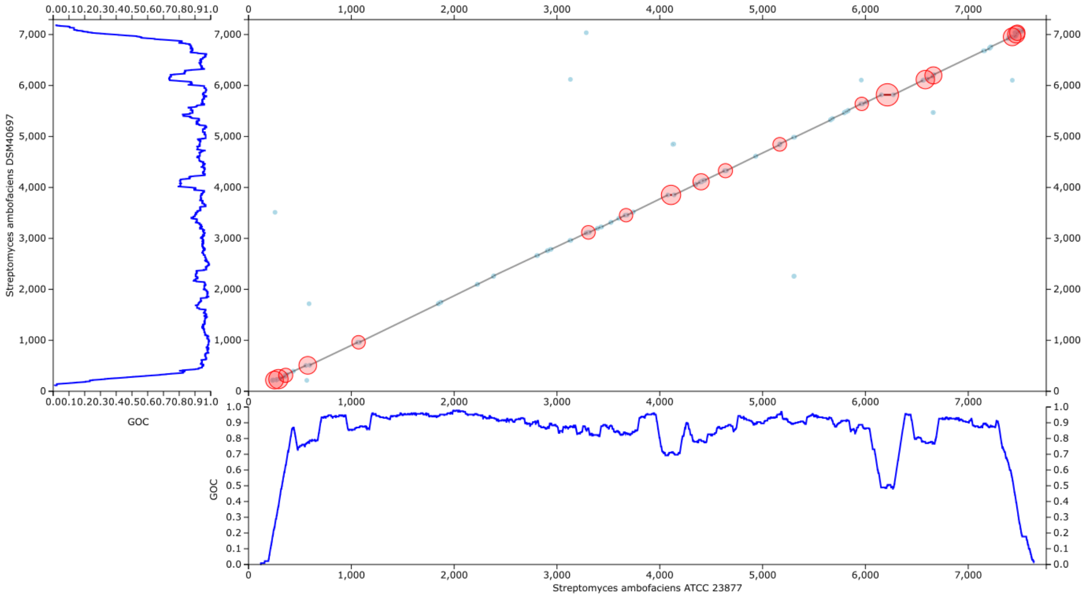
Streptomyces ambofaciens DSM40697
GOC
GOC
Streptomyces ambofaciens ATCC 23877

B

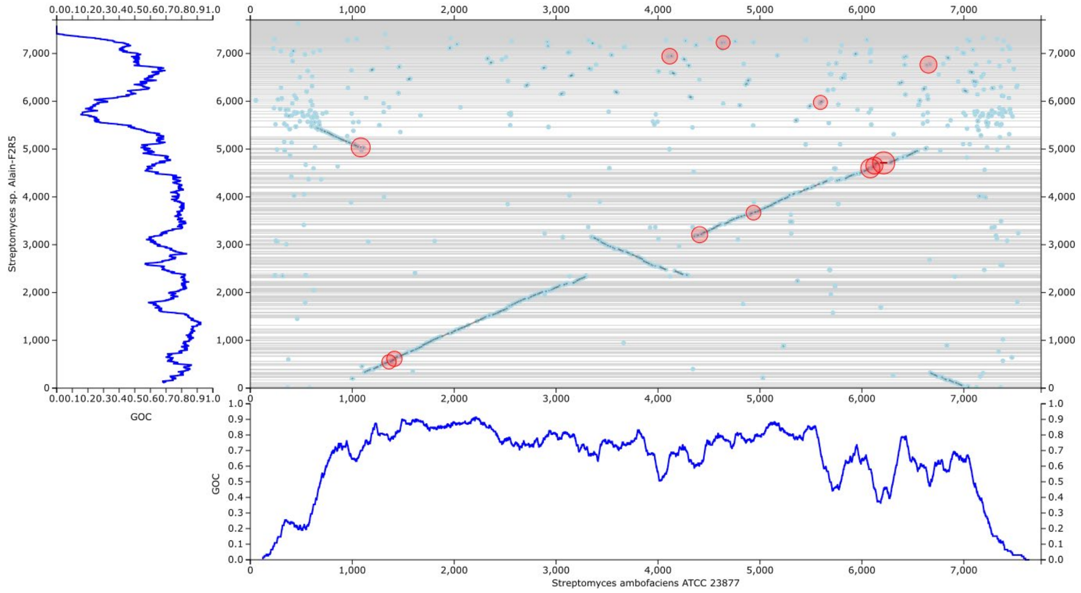
Streptomyces sp. Alain-F2R5
GOC
GOC
Streptomyces ambofaciens ATCC 23877

C

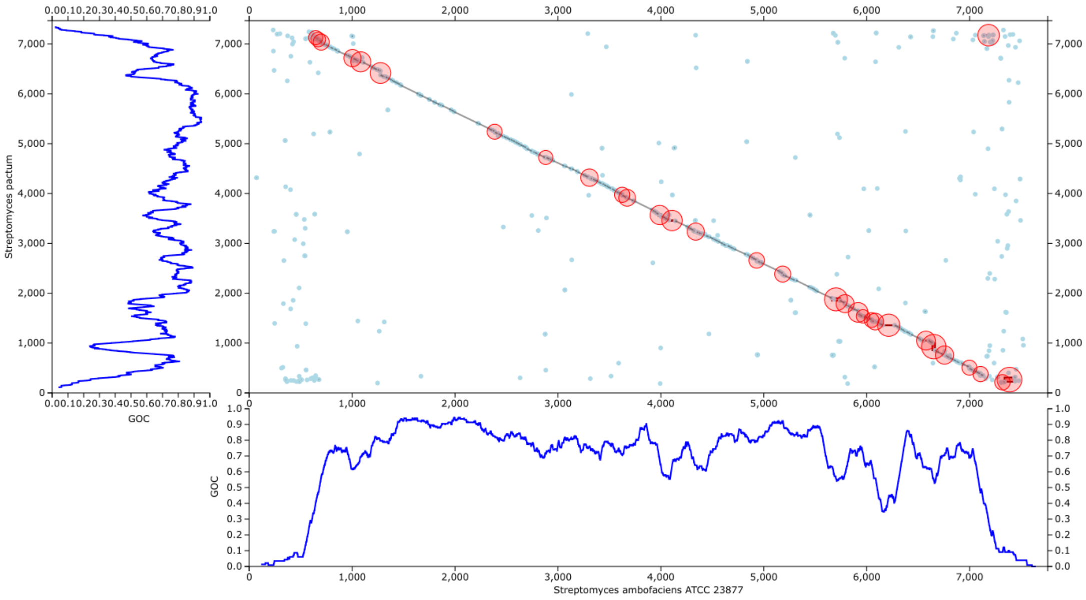
Streptomyces pactum
GOC
GOC
Streptomyces ambofaciens ATCC 23877

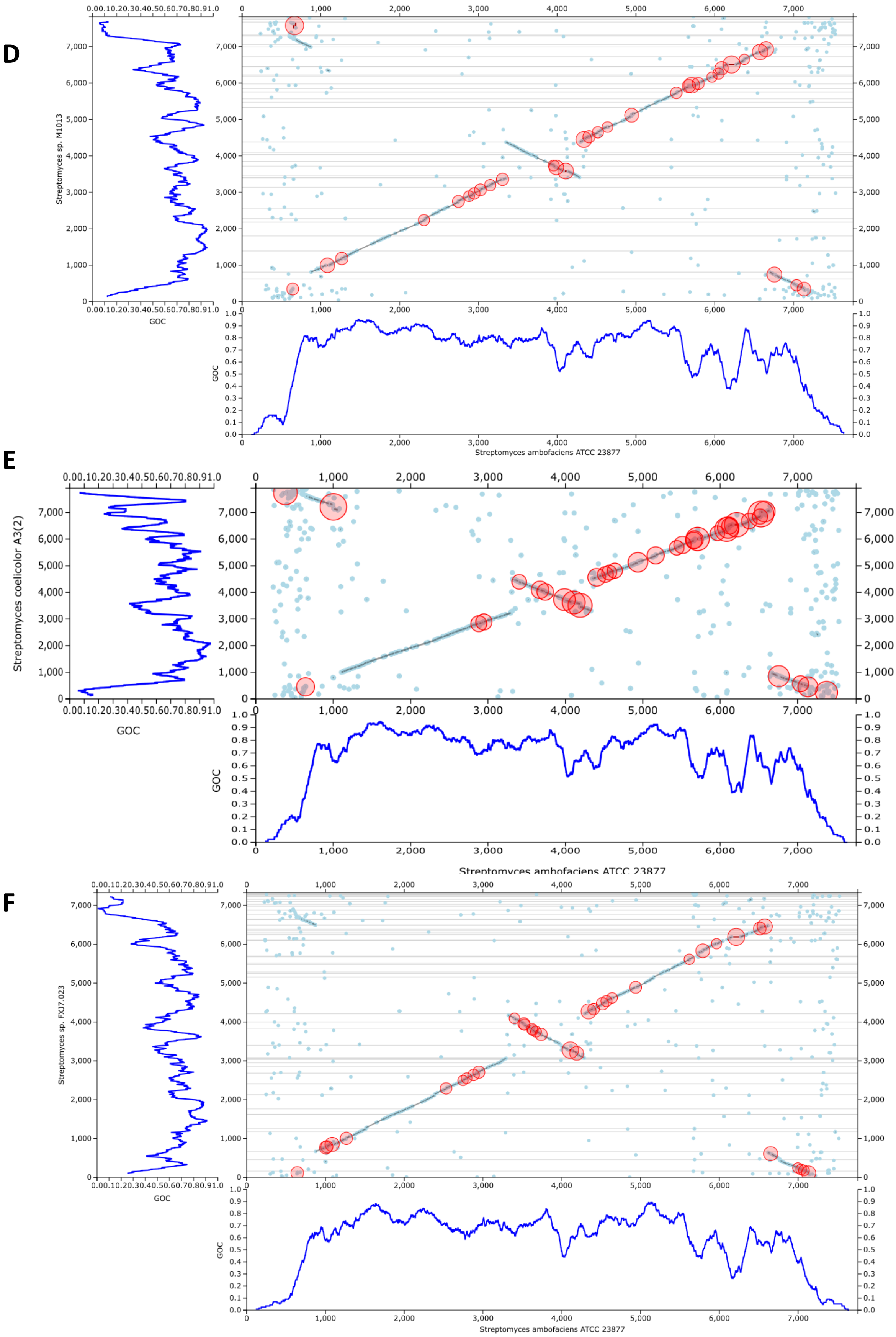
D
Streptomyces sp. M1013
GOC
Streptomyces ambofaciens ATCC 23877
E
Streptomyces coelicolor A3(2)
GOC
Streptomyces ambofaciens ATCC 23877
F
Streptomyces sp. FXJ7.023
GOC
Streptomyces ambofaciens ATCC 23877

G

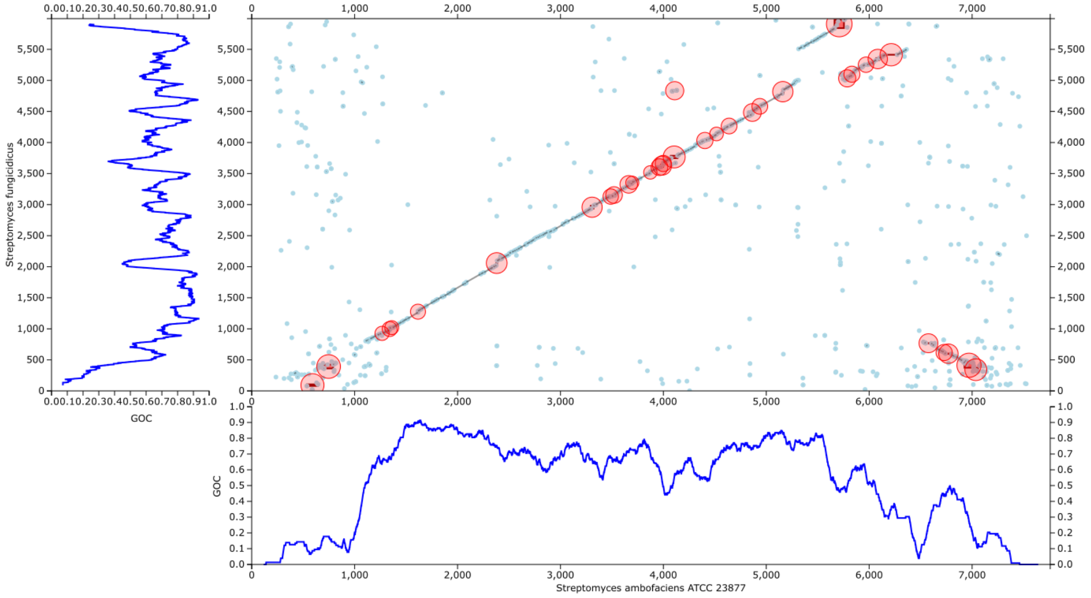

Streptomyces fungicidicus
GOC
Streptomyces ambofaciens ATCC 23877


H

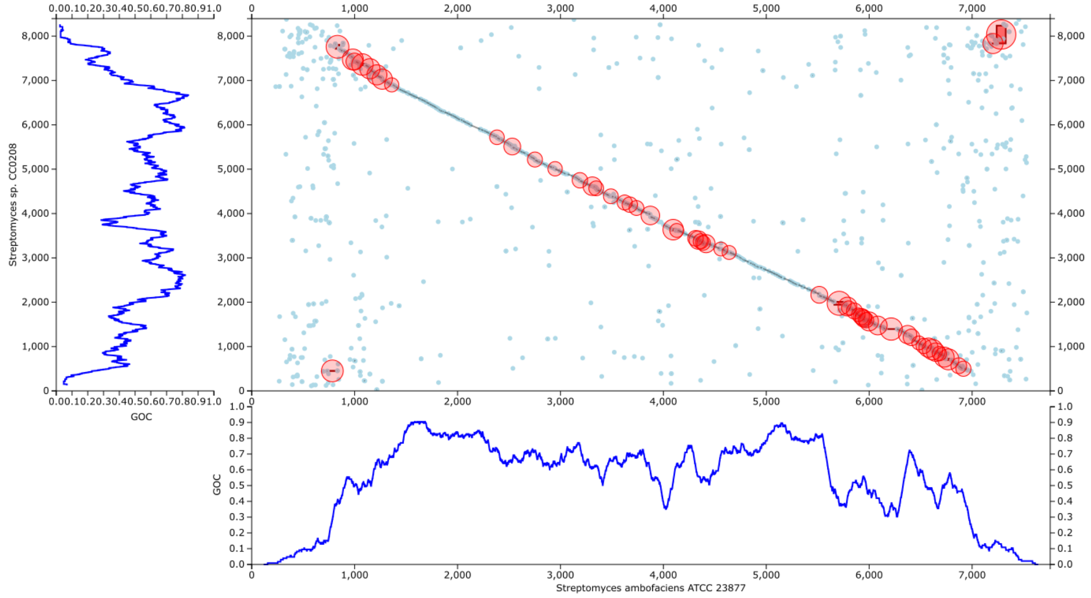

Streptomyces sp. CC0208
GOC
Streptomyces ambofaciens ATCC 23877


I

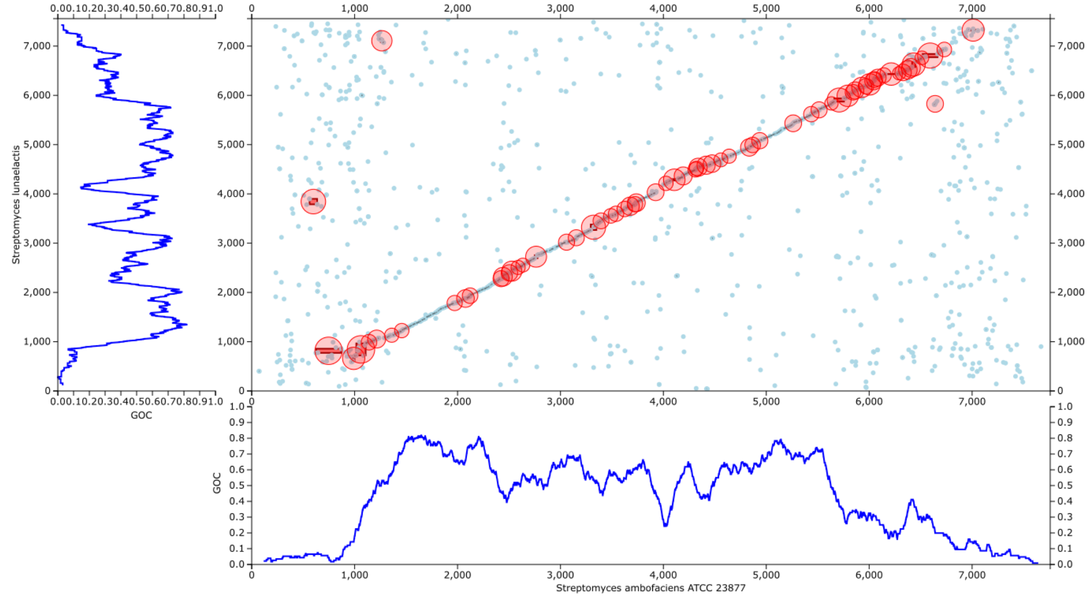

Streptomyces lunaelactis
GOC
Streptomyces ambofaciens ATCC 23877

J

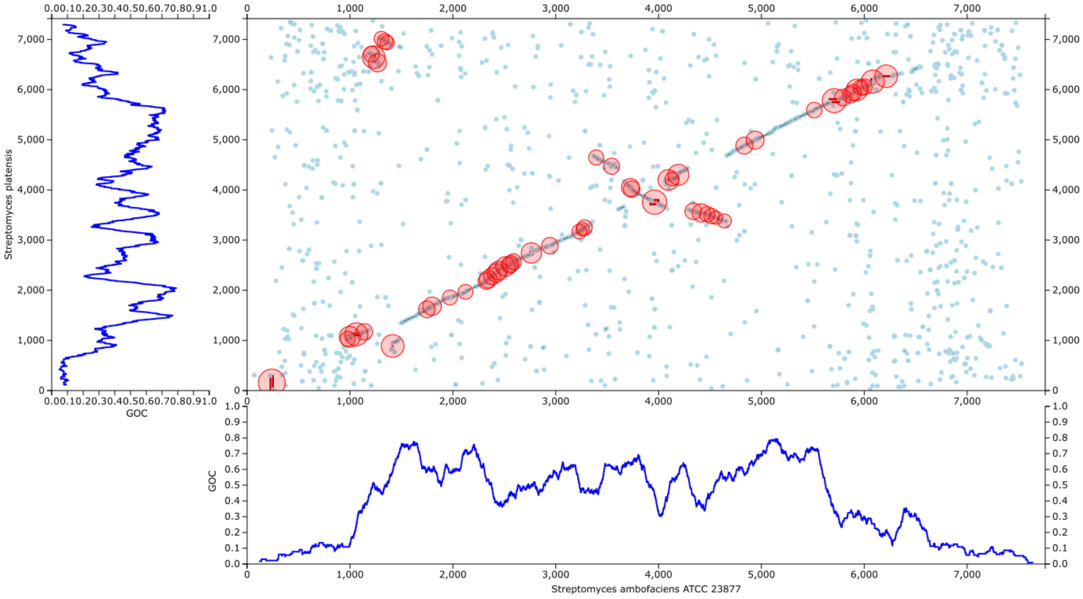
Streptomyces platensis
GOC
Streptomyces ambofaciens ATCC 23877

K

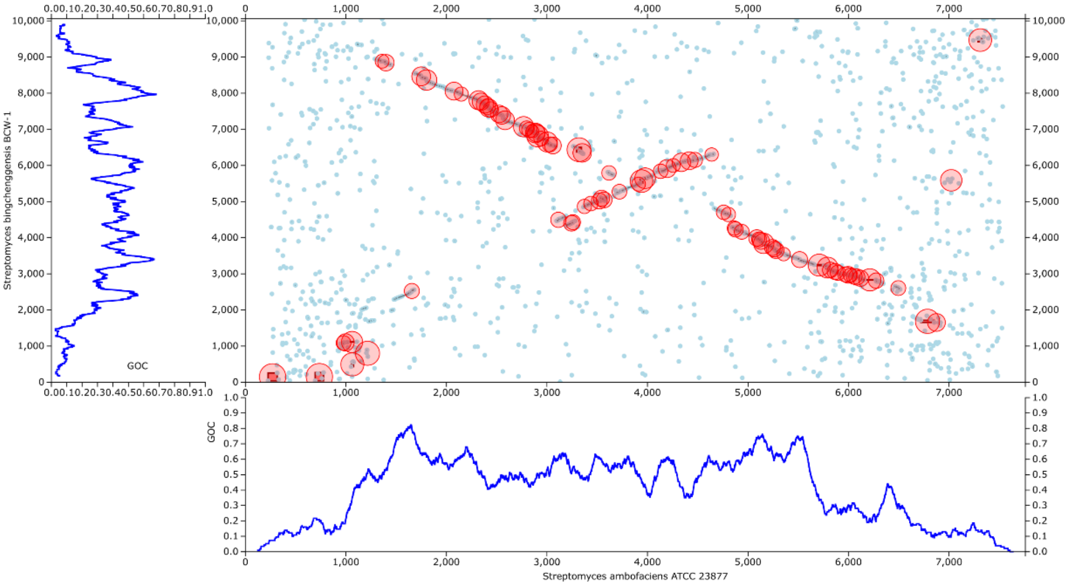
Streptomyces bingchenggensis BCW-1
GOC
Streptomyces ambofaciens ATCC 23877

L

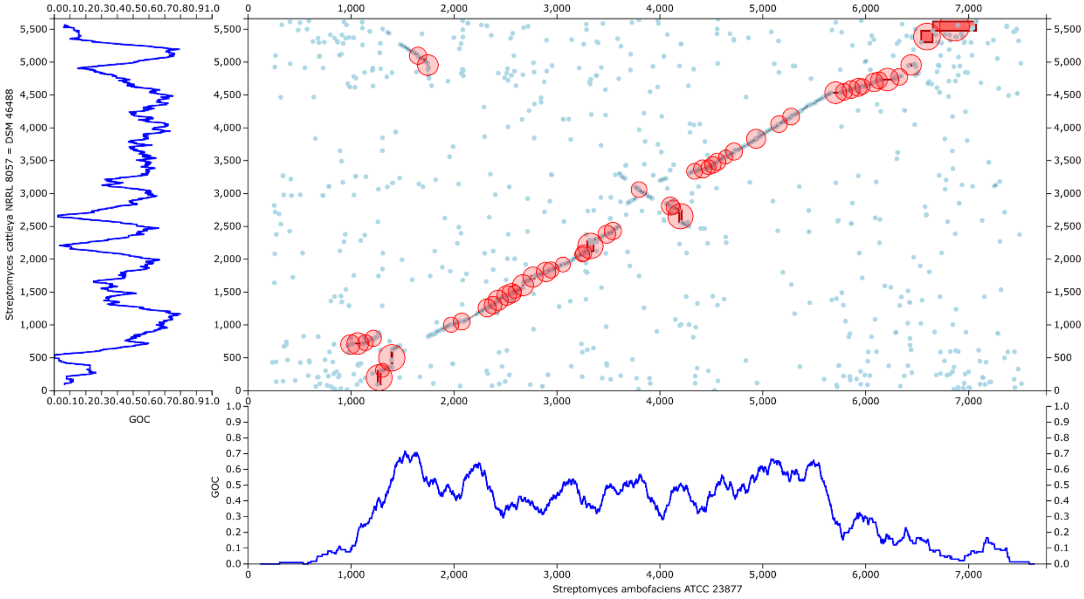
Streptomyces cattleya NRRL 8057 = DSM 46488
GOC
Streptomyces ambofaciens ATCC 23877

**Supplementary Figure S7**: Example of a genomic island (cd1336) obtained by SYNTERUPTOR, delimited by synteny regions constituted of two or three CDSs in an otherwise large non-syntenic region. A) Dotplot and GOC graphs of the chromosomes of *S. ambofaciens* ATCC 23877 and *S. coelicolor* A(3)2 showing the considered genomic island circled in yellow. B) Content of the genomic islands in *S. ambofaciens* ATCC 23877 and *S. coelicolor* A(3)2 and C) schematic representation of the *S. ambofaciens* ATCC 23877 and *S. coelicolor* A(3)2 chromosomal regions showing that genomic islands (represented in dark blue in *S. ambofaciens* ATCC 23877 and light blue in *S. coelicolor* A(3)2) are delimited by very small syntenic regions constituted in the left of three CDSs and in the right of two CDSs (in yellow) in large non-syntenic regions (dark and light grey).

**A)**

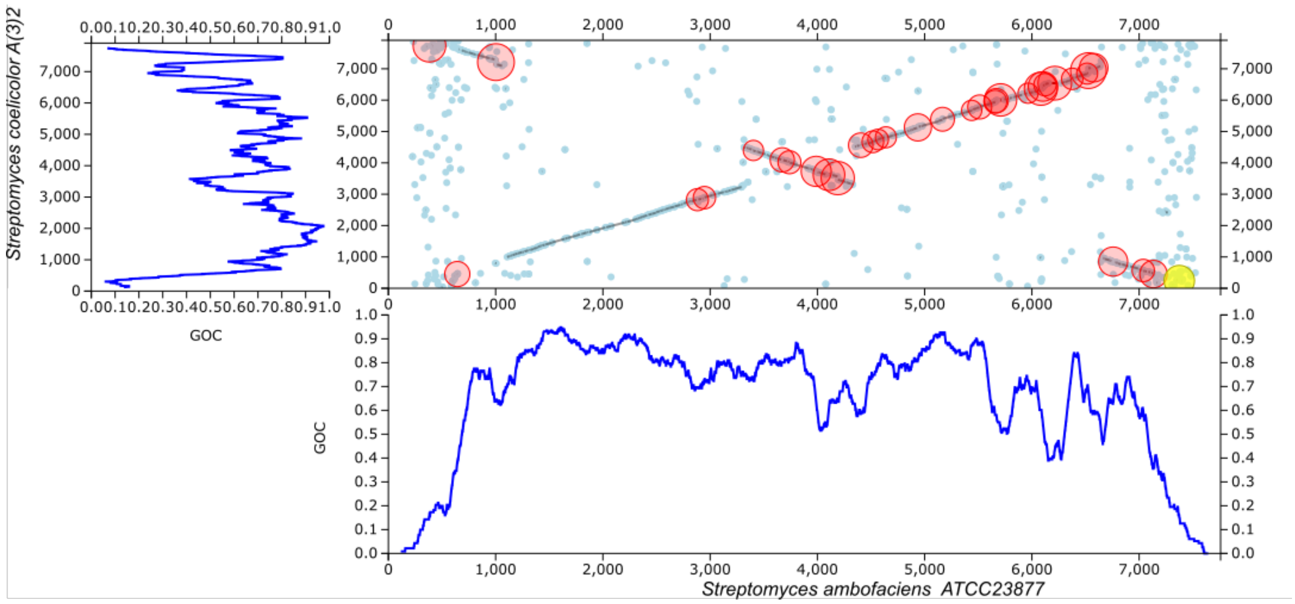


**B)**

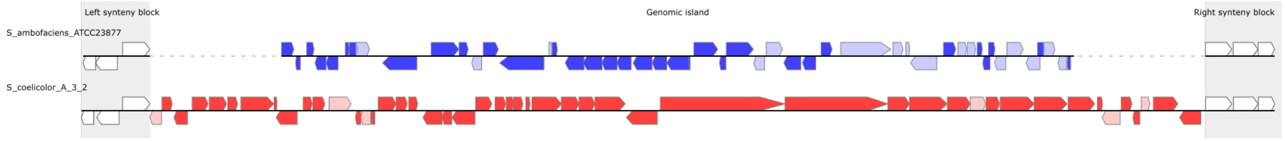


**C)**

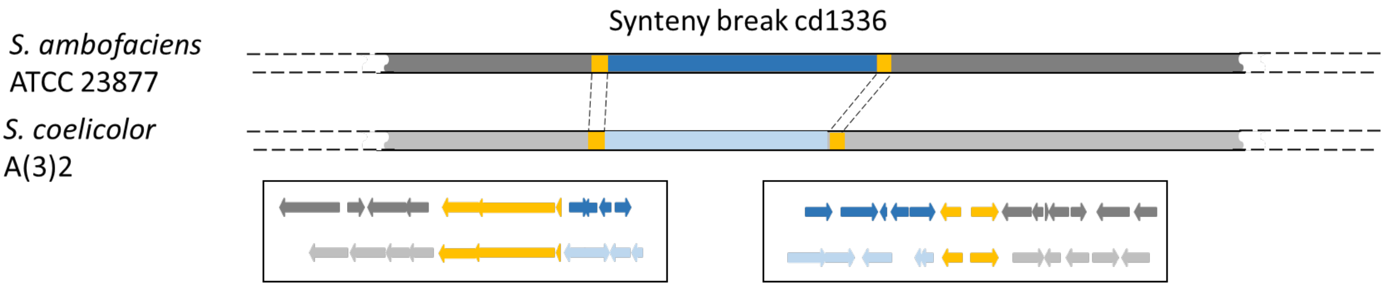

**Supplementary Figure S8**: Visualisation of Region 20 of *S. ambofaciens* ATCC 23877 genome predicted to contain four SMBGCs by antiSMASH. Red box: GI#819 detected by SYNTERUPTOR and containing the sphydrofuran SMBGC. CC: candidate cluster.

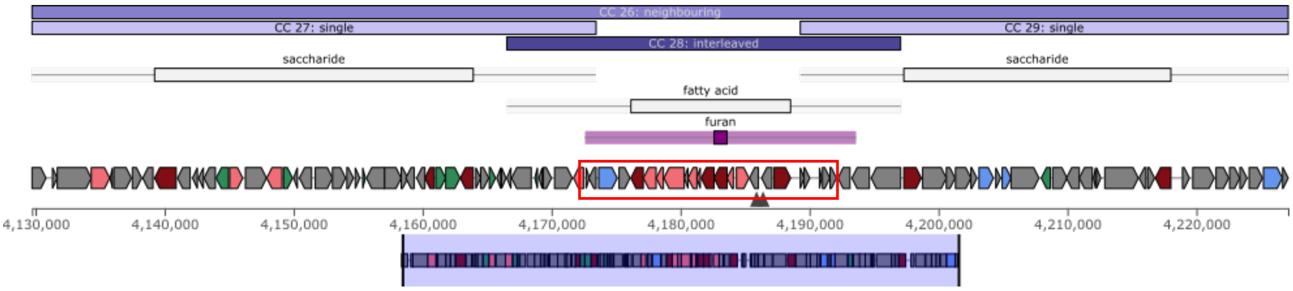
